# A mathematical model for cell-induced gel contraction incorporating osmotic effects

**DOI:** 10.1101/2021.12.08.471846

**Authors:** J. R. Reoch, Y. M. Stokes, J. E. F. Green

## Abstract

Biological tissues are composed of cells surrounded by the extracellular matrix (ECM). The ECM can be thought of as a fibrous polymer network, acting as a natural scaffolding to provide mechanical support to the cells. Reciprocal mechanical and chemical interactions between the cells and the ECM are crucial in regulating the development of tissues and maintaining their functionality. Hence, to maintain *in vivo*-like behaviour when cells are cultured *in vitro*, they are often seeded in a gel, which aims to mimic the ECM. In this paper, we present a mathematical model that incorporate cell-gel interactions together with osmotic pressure to study the mechanical behaviour of biological gels. In particular, we consider an experiment where cells are seeded within a gel, which gradually compacts due to forces exerted on it by the cells. Adopting a one-dimensional Cartesian geometry for simplicity, we use a combination of analytical techniques and numerical simulations to investigate how cell traction forces interact with osmotic effects (which can lead to either gel swelling or contraction depending on the gel’s composition). Our results show that a number of qualitatively different behaviours are possible, depending on the composition of the gel (i.e. the chemical potentials) and the strength of the cell traction forces. We observe an unusual case where the gel oscillates between swelling and contraction. We also consider on how physical parameters like drag and viscosity affect the manner in which the gel evolves.

## 1 Introduction

Biological tissues are composed of cells living in extra-cellular matrix, hereafter designated ECM (Iordan *et al*., 2010). The ECM provides mechanical support to the cells *in vivo* and helps to regulate cell behaviour, as well as playing a key role in the mechanical behaviour of the tissues themselves (Humphrey *et al*., 2014; Dolega *et al*., 2021). The ECM *in vivo* can be thought of as a fibrous polymer network; it can consist of a number of different substances, including polysaccharides, proteoglycans and proteins. To reproduce *in vivo*-like behaviour when cells are cultured *in vitro*, they are often seeded in a gel, which aims to mimic the ECM. Since the structural protein collagen is the primary component of the ECM in many animal tissues, collagen gels are frequently used in laboratory studies, but a wide range of other natural (*e.g*. Matrigel) or synthetic (*e.g*. poly(lactic acid)) gels are also used (Wade and Burdick, 2012). Improved understanding of the mechanical behaviour of biological gels, together with cell-cell and cell-gel interactions will lead to better understanding of the development and functioning of tissues.

The mechanical characteristics of a tissue can have a powerful effect on cell behaviours such as proliferation, differentiation and cell motility; an effect that is, in fact, reciprocal, since the tissue is maintained by these cells (Rozario and DeSimone, 2010; Wade and Burdick, 2012; Humphrey *et al*., 2014). Experiments aimed at gaining more insight into the cell-ECM relationship and how each regulates and affects the other, are often conducted using cell-seeded gels grown *in vitro*, where it is often easier to control conditions and make observations. One such experiment, presented by Moon and Tranquillo (1993) studies the contraction of collagen gels under the influence of cell traction forces. The cells interact mechanically with the polymer network surrounding them, leading to this polymer network being reorganised and compacted. These processes of ECM remodelling are important in tissue growth and development and, accordingly, are important in a range of related topics such as wound healing and tissue engineering.

In the Moon and Tranquillo (1993) experiment, a microsphere of collagen gel is prepared, cells are seeded within the gel, and these cells, through the traction stresses they exert on the polymer network, compact the gel over approximately two days. Although the degree of gel contraction can provide a measure of the forces exerted by the cells, it is dependent on the specific procedures employed in their experiment (*e.g*. cell density, gel composition and size, etc.). To obtain a quantitative measure of the cell-derived forces, Moon and Tranquillo (1993) developed a mathematical model of the process based on the mechanochemical theory of Murray *et al*. (1983). This model assumed that the only forces acting on the gel were those exerted by the cells; however, gels can swell or contract in the absence of cells, for example, due to osmotic effects. Recent studies have suggested that these osmotic effects could be used to manipulate the mechanical environment of cells *in vitro* by, for example, applying a compressive force to the gel in which they reside (Monnier *et al*., 2016). Our aim in this paper is to gain more understanding of how cell and osmotic forces interact within cell-seeded gels grown *in vitro* through the development and investigation of a mathematical model. In particular, we are interested in the different qualitative outcomes that may arise (*e.g*. gel contraction, swelling or dissolution), and how these outcomes, together with the dynamics of the process, depend on factors such as the gel composition, cell seeding density, cell traction strength and interphase drag between the network and solution phases of the gel.

The mathematical model of Moon and Tranquillo (1993) and a later model of Green *et al*. (2013) treat the gel as a single material (either fluid or solid) which must be assumed to be compressible for it to be able to contract. However, many types of biological gel are largely made up of water, an incompressible fluid, bringing the appropriateness of this assumption into question. Therefore, in this paper, we investigate the possibility that gel contraction is a result of syneresis, that is, as a result of water being squeezed out of the gel. This possibility is noted in both Moon and Tranquillo (1993) and Green *et al*. (2013), but was not pursued further. Osmotically-driven movement of solution into and out of gels has been studied in the context of gel mechanics using Flory-Huggins theory. This theory is derived from statistical thermodynamics, assuming that the polymer network and solvent which make up the gel exist on a two-dimensional lattice, and considering the number of ways in which the polymer and solvent molecules can be arranged on this lattice (Kumar and Gupta, 2003). From this, the entropy and enthalpy of the mixture is derived. Using these, we can calculate the free energy of the mixture per unit volume, and thereby, the osmotic pressure (Kumar and Gupta, 2003; Winstanley *et al*., 2011). This term refers to the additional pressure that must be applied to reach an equilibrium between the gel and solvent separated by a semi-permeable membrane (Winstanley *et al*., 2011). Osmotic pressure will be included in our model through chemical potential functions; this is discussed further in Section 2.3. A number of previous studies have modelled the movement of solution into and out of gels and gel-like substances, usually in the absence of cells, for example biofilms (Cogan and Keener, 2004; Winstanley *et al*., 2011), swelling polymer gels (Keener *et al*., 2011b,a), and general polymer solutions (Doi, 2011; Doi and Onuki, 1992). In the majority of these studies, the gel is modelled as a mixture of two interacting components: a polymer network, and a fluid solvent.

The multiphase or mixture theory approach provides an obvious framework for modelling the dynamics of gel swelling and contraction, and model of this type have been developed by Keener *et al*. (2011a,b) and Mori *et al*. (2013). They consider a gel consisting of two phases, polymer network and solvent, and present mass balance and momentum balance equations for each phase. Osmotic effects are incorporated in the momentum balance equations for each phase through terms involving spatial gradients of the chemical potentials. The chemical potentials depend on the Flory-Huggins free energy, and are functions of the volume fractions of network and solvent. We aim to combine their approaches with mechanochemical theory (Murray *et al*., 1983) to produce a new mathematical model for the mechanics of cell-seeded gels which incorporates osmotic effects.

The paper is structured as follows. In Section 2, we develop our multiphase model for gel-solvent interaction. This model includes the novel addition of cell-gel mechanical interactions, considering the cell traction stresses acting on the polymer network, alongside the osmotic effects introduced by free energy. As done by Keener *et al*. (2011a,b) and Mori *et al*. (2013) we non-dimensionalise the model and transform it into 1D Cartesian coordinates to obtain a simple model for analysis. In Section 4, the equilibrium conditions for the model are presented, as well as a short time solution. This short time solution allows us to evaluate how the model evolves away from steady-states and initial conditions and, so, to determine the stability of steady states. We implement the 1D model numerically in Section 5 to study the conditions under which swelling or contraction occur. Finally, in Section 6 we discuss the key results from this model and possible extensions for future investigation.

## 2 Mathematical model

We consider a gel seeded with cells, which sits within a surrounding bath of liquid solvent (*e.g*. nutrient medium). This gel-solvent system is sketched in Fig. 1. The domain Ω is divided into two regions: the gel region Ω_*g*_, and the surrounding region of pure solvent Ω_*s*_. We note that the problem can be studied in different geometries (*i.e. Ω_*g*_* need not be spherical). We let ***x*** denote position in Ω and *t* denote time. The centroid of the gel is at ***x*** = 0 and the gel-solvent interface, denoted Γ_*g*_(***x**, t*) = 0, is the boundary between Ω_*g*_ and Ω_*s*_. This interface between the gel and surrounding solvent can move over time with the movement of solvent between the two regions.

**Figure 1:**
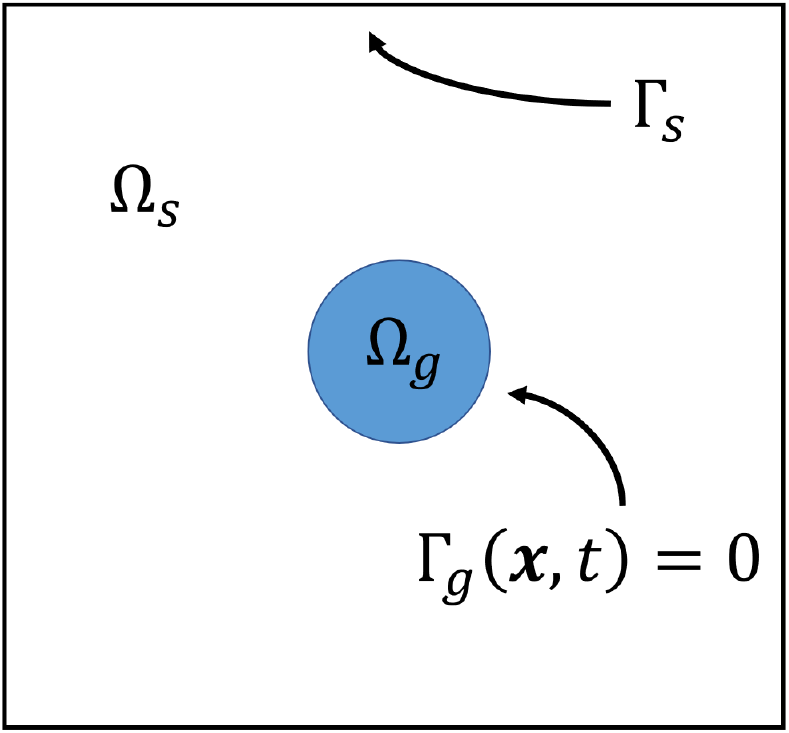
Gel-solvent domain Ω = Ω_*g*_ ∪ Ω_*s*_. Ω_*g*_ contains the cell population as well as positive volume fractions for both the polymer network and solvent, whereas Ω_*s*_ contains only solvent. Γ_*g*_(***x**, t*) = 0 is the moving boundary of the gel, also referred to as the gel-solvent interface. Γ_*s*_ is the external boundary of the domain.

We use a multiphase modelling approach based on the work of Keener *et al*. (2011b) and Mori *et al*. (2013), modified to incorporate cells. The gel is assumed to be made up of two phases, polymer and solvent, each of constant density, with volume fractions denoted by *θ_p_*(***x**, t*) and *θ_s_*(***x**, t*) respectively. Hence, we define Ω_*g*_ to be the region where *θ_p_* > 0 and *θ_s_* > 0, and Ω_*s*_ to be that where *θ_p_* = 0 and *θ_s_* = 1. Cells are only present in the gel region Ω_*g*_, and for simplicity, we assume that the volume they occupy within the gel is negligible; we therefore do not include a cell volume fraction and instead consider cell density *n*(***x**, t*), where *n*(***x**, t*) = 0 in Ω_*s*_ (similar to Barocas and Tranquillo (1994)). Thus, the no-voids condition

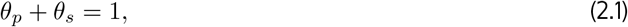

is satisfied everywhere in the domain Ω = Ω_*g*_ ∪ Ω_*s*_. Moreover, the model presented below, while written for the gel region Ω_*g*_, is also applicable to the solvent region Ω_*s*_ on setting *θ_p_* = *n* = 0 and *θ_s_* = 1.

### 2.1 Mass and momentum conservation equations

We assume the mass of both polymer and solvent is conserved (*i.e*. production and degradation of both species is neglected), so that

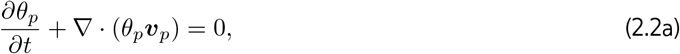

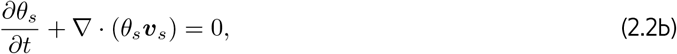

where ***v**_p_*(***x**, t*) and ***v**_s_*(***x**, t*) are the polymer and solvent velocities respectively. Given the no-voids condition (2.1), adding equations (2.2a) and (2.2b) yields

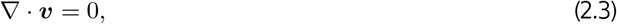

where ***v*** = *θ_p_**v**_p_* + *θ_s_**v**_s_* is the volume-averaged velocity of the polymer-solvent mixture (Keener *et al*., 2011b); we choose to replace (2.2b) with (2.3). Note that in the solvent region Ω_*s*_ we simply have ∇ · ***v**_s_* = 0.

For simplicity, cell proliferation and death are neglected, so the cell population is fixed. Cells are assumed to move by a combination of advection with the polymer network and unbiased random motion, modelled by Fick’s Law with diffusion coefficient *D*. Conservation of cells then gives

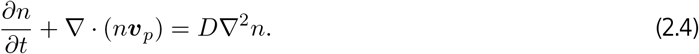

We obtain equations for ***v**_p_*(***x**, t*) and ***v**_s_*(***x**, t*) by considering the momentum balance throughout the domain. Green *et al*. (2013) note that the Deborah number (which gives the ratio of elastic to viscous effects) found experimentally for gels like collagen is small 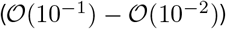, meaning that elastic effects can be ignored. Hence, following Keener *et al*. (2011b) and Mori *et al*. (2013), we model both the polymer and the solvent phases as viscous fluids with a common pressure, *P*. The viscous stresses in the two phases are encapsulated by the deviatoric tensors ***σ**_p_* and ***σ**_s_*, where the polymer stress tensor ***σ**_p_* and the rate of strain tensor ***e**_p_* are defined by

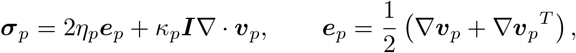

with the solvent tensors ***σ**_s_* and ***e**_s_* similarly defined (with subscript *s* in place of *p*). The constants *η_i_* and *κ_i_* (*i* = *p, s*) are the dynamic and bulk viscosities of each phase *i* respectively, and ***I*** is the identity tensor.

As in Keener *et al*. (2011b), we assume that the forces exerted on the two phases come from inter-phase drag (which is proportional to the product of the volume fractions of the two phases) and chemical potential gradients. In addition, we include traction stresses exerted by cells on the polymer network. Inertia can be neglected on the time and length scales typical of experiments such as Moon and Tranquillo (1993), so that the momentum balances for the two phases are given by

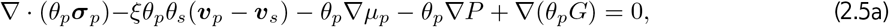

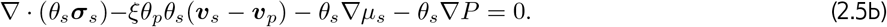

In equations (2.5a) and (2.5b), *μ_p_*(*θ_p_*) and *μ_s_*(*θ_p_*) are the chemical potentials for the polymer and solvent respectively, while *ξ* is the constant drag coefficient. The traction force exerted by the cells on the polymer network is a novel addition in this context of multiphase gel modelling; it is incorporated as a body force acting on the gel and is given by the gradient of *θ_p_G*(*n*), where *G* is a scalar potential energy function (Mori *et al*., 2013). The forms of the cell potential function *G* and the chemical potentials *μ_p_* and *μ_s_* are detailed in Section 2.2 and 2.3 below.

### 2.2 Cell potential energy function

We assume the energy potential associated with the cells to be given by the Hill equation

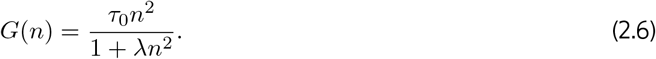

This differs from the function described in previous works (Murray, 2001; Moon and Tranquillo, 1993; Green *et al*., 2013) in having *n*^2^ rather than *n* in the numerator. This means that *∂G/∂n* > 0 for all *n*, which ensures that the cell traction force,

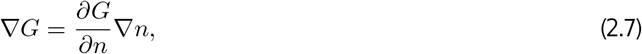

acts in the direction of increasing cell concentration. The positive parameter *τ*_0_ provides a measure for the strength of cell traction forces, and *λ* is a positive contact inhibition parameter, which reflects that the force exerted by cells decreases as the cells become more densely populated.

### 2.3 Chemical potentials

The chemical potential functions *μ_p_* and *μ_s_* describe the work done by the free energy in the polymer and solvent to affect the swelling or compaction of the gel. These are defined as

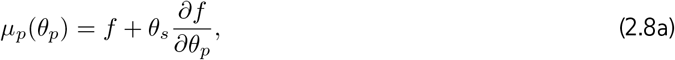

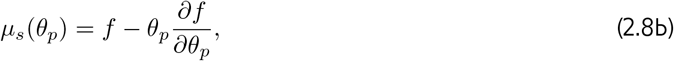

where *f*(*θ_p_*) is the free energy per unit volume of gel (Keener *et al*., 2011b). The free energy function, derived from polymer physics, is defined below. The polymer chemical potential *μ_p_* describes the change in free energy resulting from an additional polymer unit being added to the gel, while the solvent chemical potential *μ_s_* describes the change in free energy from an additional solvent unit being added (Keener *et al*., 2011b).

The free energy of the system per unit volume of gel is

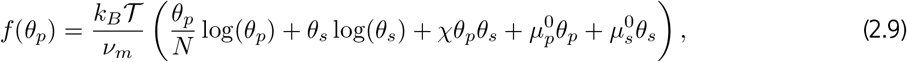

where *k_B_* is the Boltzmann constant, 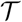 is temperature, *ν_m_* is the characteristic volume of a monomer in our system, *N* is the chain length of the polymer, *χ* is the Flory interaction parameter and the constants 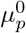 and 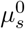 are dimensionless quantities known as the standard free energies of the polymer and solvent respectively. The logarithmic terms in the function describe the entropy of mixing polymer and solvent; these terms always encourage swelling in the gel. The latter terms involving *χ*, 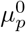 and 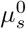 can increase the tendency for the gel to swell or contract depending on the signs of these parameters. The *χ* term describes the energy of mixing, while the terms involving 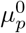 and 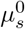 describe the interaction energy in a pure polymer or solvent state respectively (Rubinstein *et al*., 2003).

In most of the relevant literature (*e.g*. Mori *et al*. (2013); Rubinstein *et al*. (2003); Zhang *et al*. (2008)) the standard free energy parameters 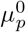 and 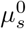 are not included, so that the free energy function represents only the interaction or mixing of the phases as opposed to the total free energy (Keener *et al*., 2011a). However, following the work of Keener *et al*. (see Keener *et al*. (2011b,a); Sircar *et al*. (2013)), we retain the standard free energy terms for generality. In the framework of Mori *et al*. (2013) that we adopt, we will see that these terms do not contribute explicitly to the final model, due to cancellations of terms involving *f*(*θ_p_*) and its derivatives. They are, nevertheless, contained implicitly through the derivation of the mixing parameter *χ* (see Rubinstein *et al*. (2003) for further detail).

The Flory interaction parameter *χ* is a dimensionless parameter that characterises the nature of the interaction between the phases in the mixture: *χ* < 0 indicates attraction between the phases, and accordingly, mixing of these components being energetically advantageous; *χ* > 0 corresponds to repulsion between the polymer and solvent, resulting in the phases preferring to separate (Rubinstein *et al*., 2003).

As in Keener *et al*. (2011b), from equations (2.8a) and (2.8b), we can derive further useful relations between the chemical potentials and free energy. Firstly, we have the relation,

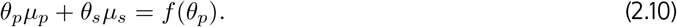

We also have that

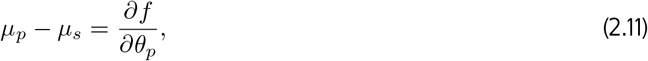

which indicates that at stationary points of *f*(*θ_p_*), the chemical potentials *μ_p_* and *μ_s_* must be equal.

### 2.4 Initial and boundary conditions

To close our system of equations, we need to impose suitable initial and boundary conditions.

The initial conditions for the volume fractions and cell density are given by

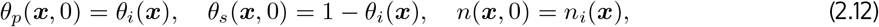

where 0 < *θ_i_*(***x***) < 1 and *n_i_*(***x***) ≥ 0 for ***x*** ∈ Ω_*p*_, and *θ_i_*(***x***) = *n_i_*(***x***) =0 for ***x*** ∈ Ω_*s*_. The initial gel-solvent interface is given by

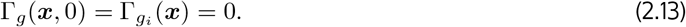

We take the centroid of the gel to be fixed in space and, therefore, have zero velocity at the origin for all time,

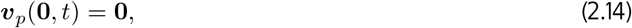

while no slip and no penetration on the external boundary of the domain Γ_*s*_ is given by

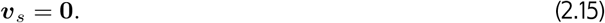

The gel-solvent interface Γ_*g*_(***x**, t*) = 0 moves over time due to movement of the polymer phase, so that its position is given by the kinematic condition

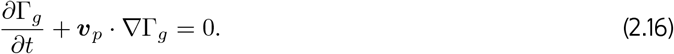

We assume there is no diffusive flux of cells out of the gel at the interface, so that

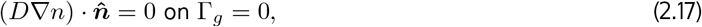

where 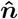 is the unit normal vector on Γ_*g*_ = 0. Continuity of stress across Γ_*g*_ = 0 is described by

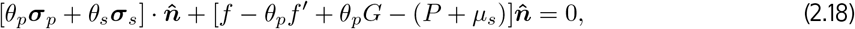

where the prime denotes differentiation with respect to *θ_p_* (Mori *et al*., 2013). The bracket notation [*J*] in equation (2.18) denotes the jump in the function *J* across the boundary; thus [*J*] = 0 indicates that *J* is continuous across the boundary. Using equation (2.8b), we simplify interface condition (2.18) to

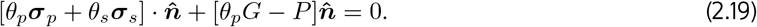

Finally, at the interface, we have

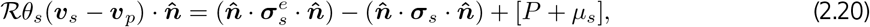

where we have introduced the superscript e to clearly designate a quantity in the solvent domain Ω_*s*_ external to the gel. This condition describes how the difference in pressure, chemical potential, and solvent stress across the interface drives fluid flow into or out from the gel, at a rate proportional to the resistance 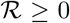 of the boundary (see equation (3.15) in Mori *et al*. (2013)). We note that an increase in 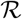 increases the resistance of the boundary so that it is more impervious to solvent flow, while a decrease indicates that it is easier for fluid to move across the boundary in and out of the gel. With the resistance 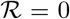, the normal solvent stresses are equal to the pressure difference across the interface.

## 3 One-dimensional Cartesian model

For simplicity, in this paper we investigate the behaviour of a one-dimensional gel (denoting the Cartesian coordinate *x*) which is assumed to be symmetrical about *x* = 0. We also assume that all quantities are continuous and differentiable at *x* = 0. Thus, we consider 0 ≤ *x* ≤ *L*(*t*) as the gel domain Ω_*g*_ in which 0 < *θ_p_*(*x, t*) < 1, *θ_s_*(*x, t*) = 1 − *θ_p_*(*x, t*), and *n*(*x, t*) ≥ 0, while the polymer and solvent velocities are *v_p_*(*x, t*) and *v_s_*(*x, t*). There is a fixed symmetry boundary at *x* = 0 and a moving boundary at *x* = *L*(*t*) (equivalently, Γ_*g*_(*x, t*) = *x* − *L*(*t*) = 0) on which the kinematic condition (2.16) becomes

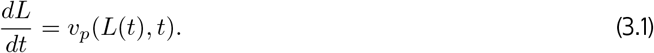

Outside the gel domain (*x* > *L*(*t*)), we have the solvent domain Ω_*s*_ with 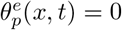 and 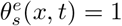, where we have introduced the superscript *e* to clearly designate the solvent domain external to the gel. From hereon, this superscript notation will be used for all quantities in Ω_*s*_, while lack of the superscript e denotes quantities in Ω_*g*_.

Since, by symmetry, *v_p_*(0, *t*) = *v_s_*(0, *t*) = 0, the continuity condition (2.3) implies that throughout Ω_*g*_ we have

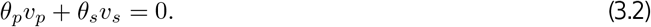

Similarly, throughout Ω_*s*_ we simply have 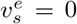, which satisfies the no-penetration condition (2.15) on the boundary Γ_*s*_. In gel domain Ω_*g*_, from the mass conservation equation (2.2a) we also have

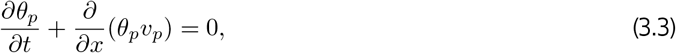

while the advection-diffusion equation for cell density (2.4) becomes

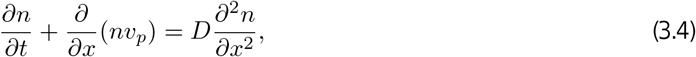

and the momentum equations (2.5a) and (2.5b) are now

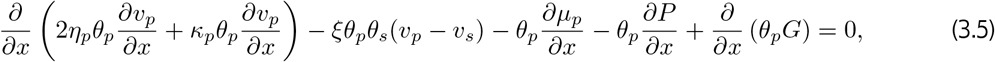

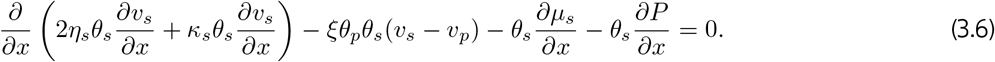

On multiplying (3.5) by *θ_s_* and (3.6) by *θ_p_* and taking the difference, we eliminate the pressure terms from the momentum equations, yielding

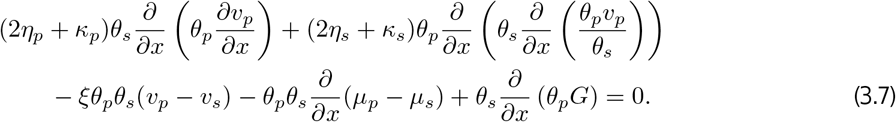

From equation (3.2), we have 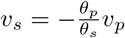, and differentiating equation (2.11) with respect to *x* gives

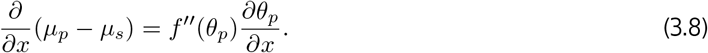

On substituting for *v_s_* and using equation (3.8), (3.7) becomes

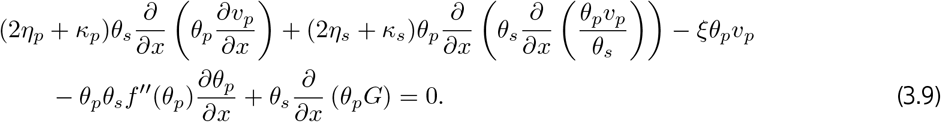

We use (3.9) to replace (3.5).

In the solvent region Ω_*s*_, where 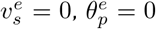 and 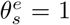, the solvent viscous stress tensor 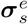 is zero, and from (2.8b) and (2.9),

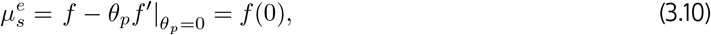

where *f*(0) is a constant. From the definition of *f*(*θ_p_*) in equation (2.9), we see that 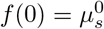. The momentum equation (2.5b) therefore simplifies to

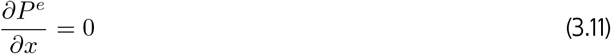

and we see that *P^e^* is at most a function of time *t*.

The 1D form of the interface condition (2.19) on *x* = *L*(*t*) becomes

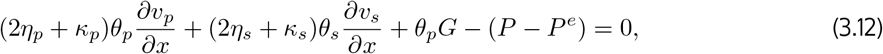

while (2.20) on *x* = *L*(*t*) becomes

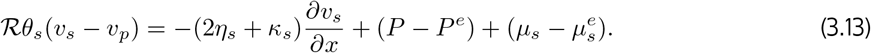

Using (3.13) to eliminate pressure from (3.12) and using (3.2) to substitute for *v_s_*, we obtain

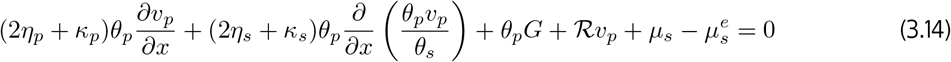

at *x* = *L*(*t*).

From (2.14), the velocity at the origin is zero for all time,

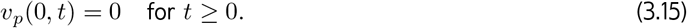

Given that *v_p_* and *∂v_p_/∂x* are continuous and differentiable at *x* = 0 and that *v_p_* is an odd function, it can be shown from equation (3.3) that if *∂θ_i_*(0)/*∂x* = 0, then *∂θ_p_*(0, *t*)/*∂x* = 0 for all *t*, *i.e*. if the polymer fraction *θ_p_* is initially symmetric and continuous about the origin, then it will remain so for all time. Since we take *θ_i_*(*x*) to be differentiable with *∂θ_i_*(0)/*∂x* = 0, we therefore have

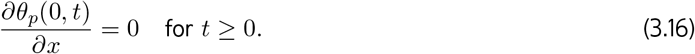

Finally, the symmetry of the cell density at *x* = 0 requires

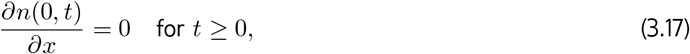

while no diffusive cell flux at *x* = *L*(*t*) requires

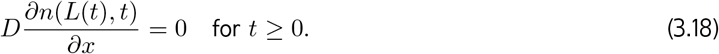

The no-voids condition (2.1), the conservation equations (3.2), (3.3), (3.4), the momentum equation (3.9), and boundary conditions (3.1), (3.14), (3.17), (3.18) comprise a complete model for the polymer and solvent volume fractions *θ_p_*(*x, t*), *θ_s_*(*x, t*), the cell density *n*(*x, t*), and the polymer and solvent velocities *v_p_*(*x, t*), *v_s_*(*x, t*) in the gel, along with the length *L*(*t*) of the gel. To solve for the pressure difference between the gel and solvent regions, *P*(*x, t*) − *P^e^*(*t*), we must add the momentum equation (3.6) and the boundary condition (3.13). However, we choose not to solve for pressure throughout the gel, given that we can study the mechanics driving the gel without its inclusion, and so drop (3.6) and (3.13) from the model.

### Non-dimensionalisation

Let 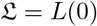 be the length scale, 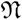 be a characteristic cell density (typically the mean initial density), and let the time scale be the ratio of polymer viscosity *η_p_* to the free energy scale 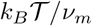. Using these scales, we non-dimensionalise our model variables as follows, where tildes denote dimensionless quantities,

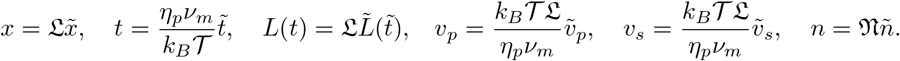

We also define the following dimensionless model parameters, again denoted by tildes,

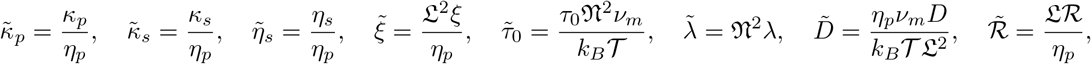

and the dimensionless free energy, chemical potential functions, and cell potential energy,

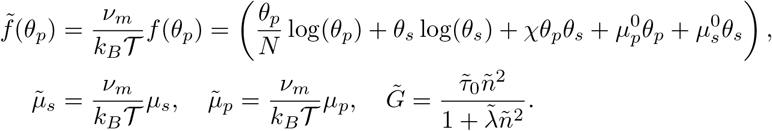

The mass balance equations (3.2), (3.3) and (3.4) are unchanged in form on writing them in terms of the scaled variables and parameters. Similarly, (2.8a), (2.8b), (2.10), (2.11) and the boundary conditions (3.1), (3.17), (3.18) are unchanged in form. Hence we do not re-write them here. The scaled forms of the momentum equation (3.9) and interface stress condition (3.14) become

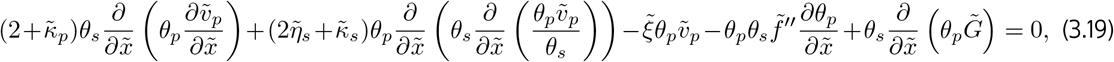

over 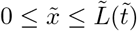 and, at 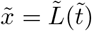,

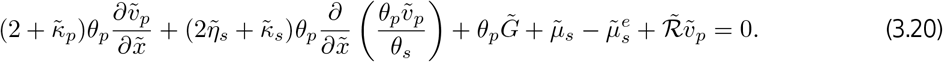

We now introduce a change in coordinates to shift our moving boundary problem onto a fixed domain. We define new coordinates 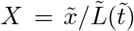 and 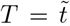, so that the domain 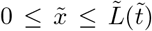 is mapped to the fixed domain 0 ≤ *X* ≤ 1. On this fixed domain the model becomes, using dots to denote differentiation with respect to time *T*, primes to denote differentiation with respect to *θ_p_*, and dropping tildes on dimensionless variables and parameters for convenience,

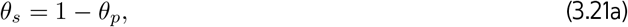

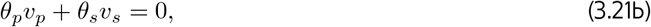

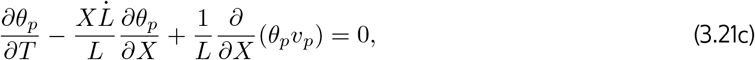

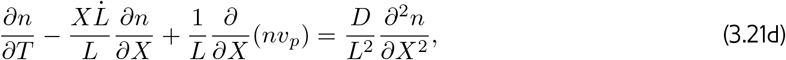

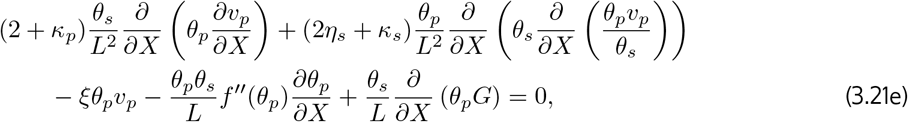

with boundary conditions at *X* = 0,

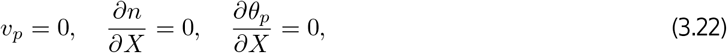

and boundary conditions at *X* = 1,

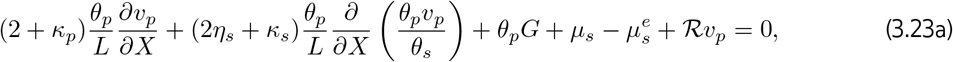

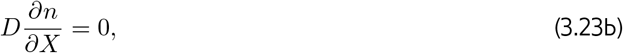

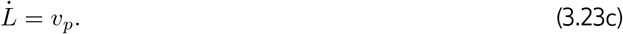

In addition, we must specify suitable initial conditions

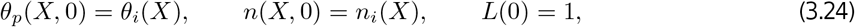

with *θ_i_*(*X*) chosen such that *∂θ_i_*(0)/*∂X* = 0. This completes our derivation of the 1D model.

## 4 Steady state and short-time behaviour

We now consider steady state (*i.e*. long time) and short time solutions of our model. The former allows us to understand the necessary conditions for the gel to equilibrate. The latter provides insight into the stability of steady states, and helps us to verify our numerical simulation methods.

### 4.1 Steady state conditions

The system reaches equilibrium when *θ_p_* and *n* are such that there is zero net force everywhere, the velocities of polymer and solvent are zero everywhere, and 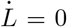, *i.e*. the free boundary stops moving. We find that steady state solutions which are spatially uniform and non-uniform in *θ_p_* and *n* are both possible.

At a steady state, the mass and momentum conservation equations (3.21c) - (3.21e) are

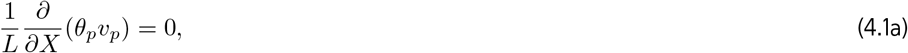

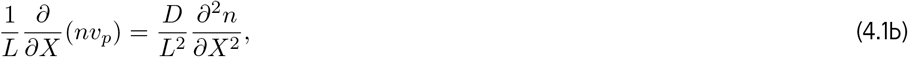

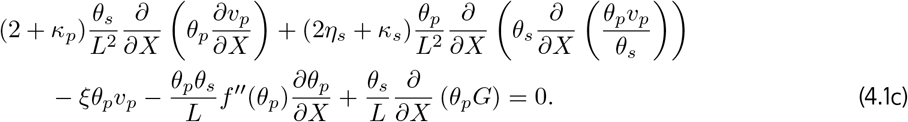

To demonstrate that the velocities are zero at equilibrium, we integrate the steady state polymer advection equation (4.1a) with respect to *X* and apply the boundary condition *v_p_* = 0 at *X* = 0. This gives *θ_p_v_p_* = 0, and since we must have *θ_p_* > 0, we see that *v_p_* = 0 must hold at equilibrium. Accordingly, we must also have *v_s_* = 0 using equation (3.21b).

The momentum balance equation (4.1c) then yields the following equilibrium condition within the gel,

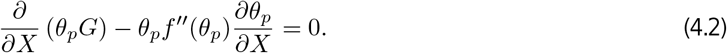

From (2.8b), we note that

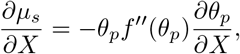

and accordingly, equation (4.2) can be expressed in the form

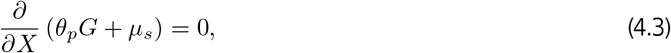

*i.e*. for the gel to be in equilibrium, the cell traction force must be balanced by the force due to chemical potential gradients. We note that with *n* = 0 (and hence *G* = 0), this is the same condition as in Keener *et al*. (2011b). Equation (4.3) is subject to the condition (3.23a) at the interface *X* = 1, which, at equilibrium, gives

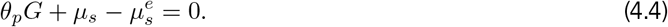

After integrating (4.3) and using (4.4) to set the constant of integration, we obtain

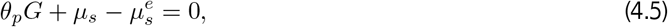

which applies everywhere at equilibrium. We note that, as shown in (3.10), 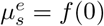 is constant.

From the cell advection-diffusion equation (4.1b), we see that, at equilibrium, we must have

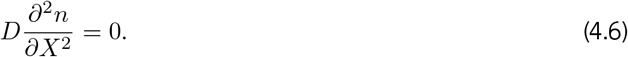

For *D* = 0, this is trivially satisfied, and equation (4.5) is sufficient for the gel to equilibrate. In this case, it is possible to have equilibrium solutions where *θ_p_* and *n* depend on *X*. With *D* ≠ 0, after integrating (4.6) and applying the no-flux condition (3.23b), we find that

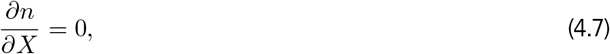

*i.e*. at equilibrium, *n* must be spatially uniform.

Now, given spatially uniform *n*, equation (4.2) can be written

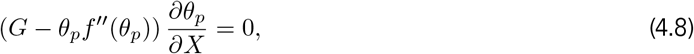

indicating that either *G* − *θ_p_f*′′ or *∂θ_p_*/*∂X* must equal zero for equilibrium. Given the functional form of *f*(*θ_p_*) (see (2.9)), we have

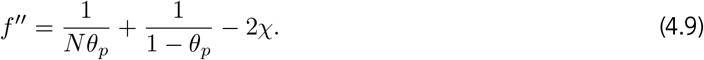

Evaluating *G* − *θ_p_f*′′ = 0, we find the following quadratic expression in *θ_p_*,

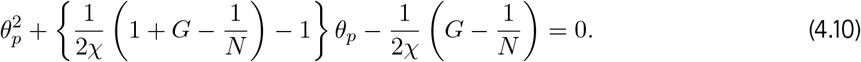

This shows that, at most, we can have two positive solutions for *θ_p_*, depending on the values of the model parameters. However, given that *θ_p_* must be continuous, only a constant value of *θ_p_* will satisfy this condition. Thus, we must have *∂θ_p_*/*∂X* = 0 to satisfy *G* − *θ_p_f*′′ = 0 at equilibrium. Therefore, from equation (4.8), we see that we must have at equilibrium,

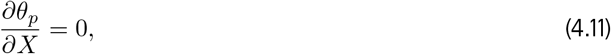

*i.e*. spatially uniform *θ_p_*. Therefore, if diffusion *D* ≠ 0, *n* and *θ_p_* must be spatially uniform and satisfy equation (4.5) for the gel to reach a steady state.

We note that, since the total mass of polymer and cells is conserved, any changes to the initial polymer fraction or cell density that change the total polymer or cell mass will lead to changes in *L*, *θ_p_* and *n* at later points in time. Hence, the equilibrium solutions for *θ_p_*, *n* and *L* are not independent of the initial conditions.

We can use equation (4.5) to calculate the spatially uniform equilibrium values of *θ_p_* and *n* for a given set of parameter values (note that this does not preclude the existence of spatially-varying steady states for such parameter values when *D* = 0). This is useful for two reasons: firstly, it provides a method to confirm that numerical simulations (such as we will see in Section 5) find the correct equilibrium values; secondly, it allows us to analyse how the equilibrium values of polymer and cell density change as chosen parameter values are adjusted. Together with a condition derived from the short time solutions in Section 4.2.2, we will use this in Section 4.2.3 to analyse steady state values of particular variables and parameters, as well as the stability of these equilibria as different parameter values change.

### 4.2 Short time analysis

We next study the behaviour of the system (3.21a) - (3.24) on a short time scale to investigate the early time evolution of the system from non-equilibrium initial conditions. In Section 4.2.2, we will determine how the system evolves over small time in response to small spatial perturbations to equilibria.

#### 4.2.1 Evolution from non-equilibrium initial conditions

We proceed by introducing the short time scale *δ* ≪ 1 and define 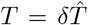. We then write the dependent variables as power series in *δ*, expanding about the initial conditions:

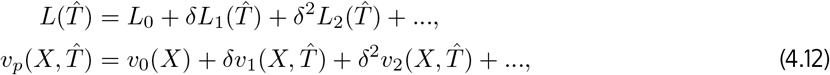

with expansions for *θ_p_* and *n* similar to that for *v_p_*. Here *L*_0_ = 1, *v*_0_(*X*), *θ*_0_(*X*), and *n*_0_(*X*) are the initial conditions. For simplicity, and in the interests of finding an analytic solution, we shall restrict our attention to spatially uniform initial conditions for *θ_p_* and *n*, *i.e*. *θ*_0_(*X*) = *θ_i_* and *n*_0_(*X*) = 1, where *θ_i_* is constant and we have scaled *n* using the initial cell density as its characteristic value.

We substitute these expansions into (3.21c) - (3.23c), and solve to obtain:

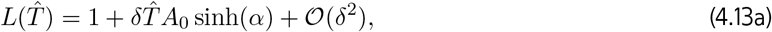

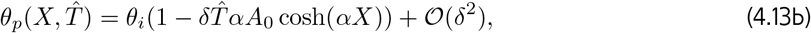

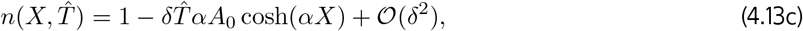

where we have introduced the parameters

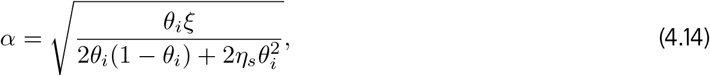

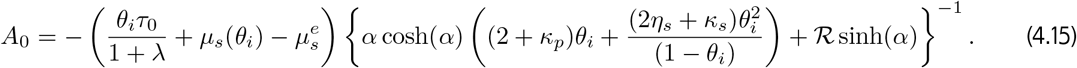

From these solutions we see that the evolution of the gel from uniform initial conditions depends on the sign of *A*_0_, which is determined by the first term in brackets in (4.15). Hence, the expansion or contraction of the gel depends on the balance between the initial cell and chemical potentials. With positive *A*_0_ (*e.g*. with a large negative mixing parameter *χ* in *μ_s_* encouraging gel swelling), the length *L* increases as the gel stretches in response to the influx of solvent, causing *θ_p_* and *n* to decrease. Conversely, for negative *A*_0_ (*e.g*. driven by a large cell traction parameter *τ*_0_), the gel contracts in length as solvent is forced out, with *θ_p_* and *n* increasing accordingly. (Note that (4.5) implies the relevant term is equal to zero if the initial condition *θ*_0_ = *θ_i_*, *n*_0_ = 1 is a steady state.) For *α* > 0 (equivalently, *ξ* > 0), the cosh(*αX*) functions in (4.13b) and (4.13c) increase monotonically with *X* and hence *θ_p_* and *n* change most rapidly at the gel’s interface. We note that, in the limit where there is no drag (*ξ* = 0, or equivalently, *α* = 0), the solutions (4.13b) and (4.13c) become spatially uniform. However, the expansion or contraction of the gel is governed by the same balance between the initial cell and chemical potentials (Reoch, 2020).

These solutions show us how the early time expansion or contraction behaviour of the gel depends on both the chemical and cell potentials when the gel is not initially in equilibrium. In the following section, we will consider how the gel behaves in response to spatial perturbations of a steady state, with the aim of gaining insight into the stability of equilibria.

#### 4.2.2 Short-time behaviour of spatial perturbations to equilibria

We now examine how the system evolves over short time from initial conditions that are small amplitude spatial perturbations to equilibrium solutions. This will suggest the stability of the equilibrium state: an equilibrium will be taken as unstable if spatial perturbations increase in amplitude over time, leading the system to evolve away from the equilibrium; an equilibrium will be taken as stable if the perturbations decay. (We note that these stability criteria are supported by our numerical solutions to the model.) For simplicity, we restrict our attention to spatially uniform equilibria as required in the general case where *D* ≠ 0.

We denote the dimensionless steady state by asterisks, *L*\*, *θ*\*, *n*\*, *v*\*, where *v*\* = 0, and *L*\* = *n*\* = 1 (since length and cell density are scaled on their equilibrium values). We take *δ* to be the short time scale as in the previous section and let *ϵ* be the amplitude of the spatial perturbation, where *δ* ≪ *ϵ* ≪ 1. Next, we take the series (4.12), *etc*., and expand each of the terms *L_j_*, *v_j_*, *θ_j_*, *n_j_*, *j* = 1, 2,…, in powers of *ϵ* with the initial conditions

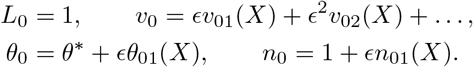

We set *θ*_01_ = cos(*γX*), *n*_01_ = *N*_01_ cos(*γX*), where *N*_01_ is an 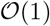 constant describing the magnitude of the spatial perturbation to the cell density. We note that higher order terms of *v*_0_ must be determined in such a way as is consistent with the other initial conditions. Note that *θ*_0_ and *n*_0_ satisfy the symmetry boundary conditions (3.22) at *X* = 0 for any choice of *γ*, while the no-flux cell boundary condition (3.23b) at *X* =1 requires that *γ* = *Zπ* for some integer *Z*. For *Z* = 0, the spatial perturbation is constant and so effectively only shifts our initial condition, resulting in a similar solution to that presented in Section 4.2.1. Furthermore, changing the sign of *Z* does not change *θ*_0_ or *n*_0_. We therefore restrict our analysis to positive values of *Z*. We also note that the condition *γ* = *Zπ* ensures that our choices of *θ*_0_ and *n*_0_ are such that the total masses of polymer and cells over the domain 0 ≤ *X* ≤ 1 are unchanged from the unperturbed initial masses (*θ*\* and 1, respectively).

Incorporating these spatial perturbations in *ϵ*, the expansions (4.12), *etc*., become

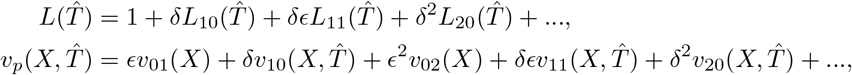

and so on. We substitute these into the governing equations to obtain solutions describing the small time behaviour. Since the calculations are standard but somewhat lengthy, the details are omitted (but can be found in Reoch (2020)). The solutions for the case with non-zero drag (*ξ* ≠ 0) are:

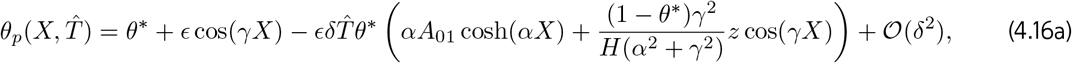

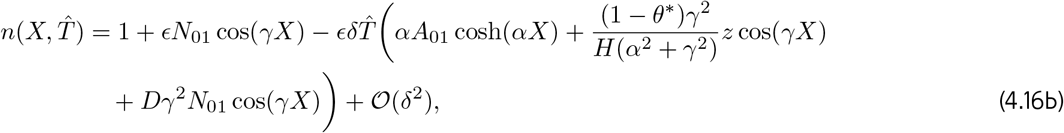

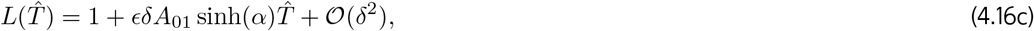

where *α* is as defined in (4.14) on replacing *θ_i_* with *θ*\*, and we have introduced the parameters

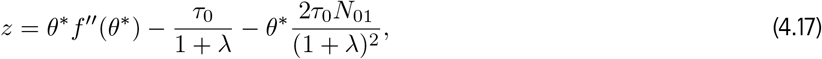

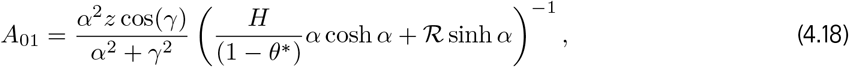

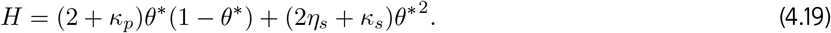

For the no drag case (*ξ* = 0) the solutions simplify to:

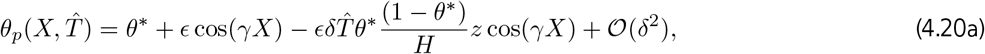

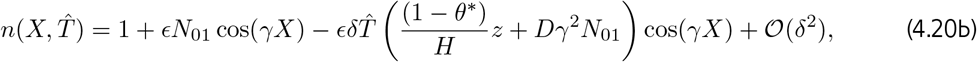

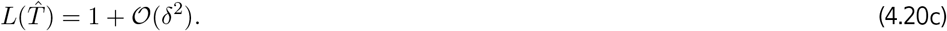

We note that, at 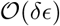, equation (4.16b) does not satisfy the no-flux cell boundary condition at *X* = 1 due to the cosh(*αX*) term. As explained in (Reoch, 2020), this is because we neglect a higher-order term involving *∂*^2^*n*_11_/*∂X*^2^, meaning that we have a singular perturbation problem and cannot satisfy all boundary conditions for *n*_11_. For the purposes of our analysis herein, we will continue to discuss this solution, as the error will be confined to the small region near *X* = 1. We also observe that in the zero-drag case, the boundary condition at *X* = 1 is, in fact, satisfied.

We see from equation (4.16c) that *L* increases or decreases depending on the sign of *A*_01_, which in turn depends on the value of *z* and the choice of *γ*. The *A*_01_ sinh(*α*) term describes the 1D expansion or contraction of the gel over the short time scale as a result of the spatial perturbation. The changes in *θ_p_* and *n* due to this expansion or contraction are described by the *A*_01_ cosh(*αX*) terms in their solutions; as previously observed, cosh(*αX*) increases monotonically with *X* since *α* > 0. Greater changes in *θ_p_* and *n* therefore occur as *X* increases across the spatial domain. This is similar to the solution given by equations (4.13a) - (4.13c), where the gel length is governed by a sinh(*α*) term, while the cosh(*αX*) terms determine the increase or decrease of the polymer fraction and cell density.

The trigonometric terms in the solutions for *θ_p_* and *n*, (4.16a) and (4.16b) respectively, describe whether the initial spatial perturbations increase in amplitude or decay over time. Growth in these perturbations is akin to an unstable equilibrium where the gel evolves away from its steady state; decaying perturbations meanwhile correspond to a stable equilibrium. In the zero-drag solution (4.20a)-(4.20c), *L* remains constant to 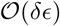 and no hyperbolic functions appear. Accordingly, *θ_p_* and *n* evolve in space with no change in the gel’s length, *i.e*. there is no flow at *X* = 1 when *ξ* = 0. Thus, a change in gel length only occurs over short time when *ξ* > 0. Since we are primarily interested here in the evolution of the polymer and cell distributions, we focus our attention on the case where *ξ* = 0. In this case, changes in the amplitude of the perturbations are simply governed by the coefficients of the trigonometric terms, avoiding any complications resulting from the small changes in gel length with drag present.

#### 4.2.3 Steady state stability conditions

We now use our results from Section 4.2.2 above to investigate the response of steady state solutions to spatial perturbations. As explained above, to simplify the interpretation of the stability results presented in this section, we restrict our attention to the case where *ξ* = 0. By considering the amplitudes of the cos *γX* terms in (4.20), we see that for the amplitude of the initial perturbations to both *θ_p_* and *n* to be decreasing in time, and accordingly, for the solution to revert back to equilibrium, we require

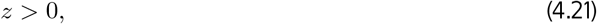

which can be expressed in the form

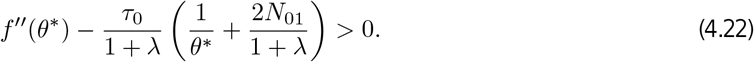

Conversely, in the case that *z* < 0, the amplitude of the perturbation to *θ_p_* will grow with time. Hence, the growth or decay of perturbations in this system is dependent on the balance between free energy and cell force. In the absence of cells, it is the sign of *f*′′(*θ*\*) which determines the stability of an equilibrium. Given that

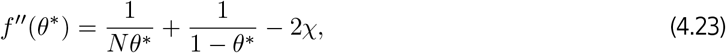

where *N* is typically large, we see that it is the sign and magnitude of the mixing parameter *χ* together with the equilibrium fraction of solvent 1 − *θ*\* that primarily determines this. Adding cells to the model will always reduce the value of *z*, and with other values held constant, move the equilibrium towards an unstable state. Similarly, with cells present, increasing cell traction strength *τ*_0_ or reducing contact inhibition *λ* will make instability more likely.

Using the stability condition (4.22), we can determine whether equilibria are stable or unstable and study how both the equilibrium and its stability change as we adjust particular parameter values. The diagrams presented in this section have been generated by solving the equilibrium condition (4.5), which, upon substituting for *G* and *μ_S_*, and recalling that we have scaled the system such that the equilibrium cell density *n*\* = 1, becomes

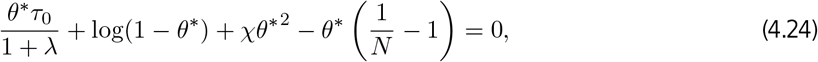

while the first term vanishes for *n*\* = 0. We solve this expression for a chosen parameter as we vary *θ*\* between 0 and 1, holding other parameters fixed. Alternatively, we find the relationship between two parameters using (4.24) while keeping *θ*\*and other parameters fixed. Through this analysis, we can also determine whether spatially uniform steady states exist for a particular set of parameter values and initial conditions.

Figs. 2a - 2c demonstrate how *θ*\* varies as we change the mixing parameter *χ* at different values of *τ*_0_ (we note that changes in the dimensionless parameter *τ*_0_ can correspond to changes in the characteristic cell density – here the physical steady state value – or the cell traction strength). In the majority of cases, larger values of *χ* indicate greater levels of contraction in the gel, corresponding to larger values of *θ*\*; this outcome should be expected, as increasing *χ* indicates that separation of the two phases in the gel is more favourable. In Fig. 2a, it is shown that in the absence of cells (n^*^ = 0), two equilibrium values of *θ*\* – one stable and one unstable – exist for the same parameter values. Note that the stability of these steady states is determined using equation (4.22). We also see that, in the absence of cells, steady states *θ*\* exist only for positive values of *χ* (given *N* = 100); with *χ* < 0, the terms in the free energy function all promote mixing between solvent and polymer, and accordingly, the gel keeps expanding until it dissolves. In this example, if a gel’s initial fraction of polymer *θ_i_* is in the region beneath the blue solid line, the gel will contract to equilibrium with a greater value of *θ_p_*, while if *θ_i_* is above the branch of stable equilibria, the gel will swell to a steady state. For *χ* < 0.62, there are no steady states possible for any initial condition *θ_i_* (*i.e*. the gel dissolves). This indicates that small changes to the initial composition of a gel, *e.g*. the fraction of polymer or make-up of the solvent, could have significant impacts on its subsequent behaviour and possible steady state.

**Figure 2:**
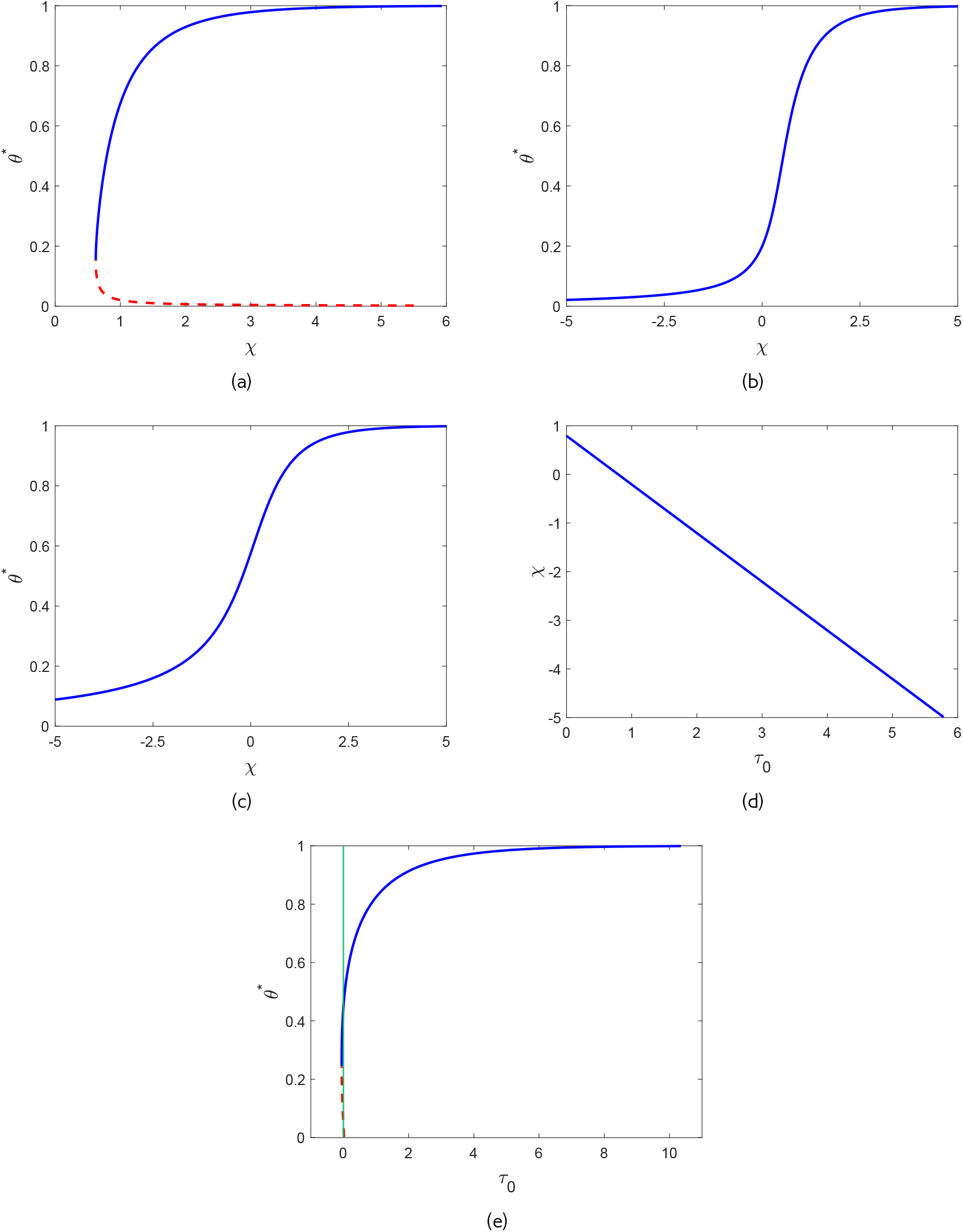
Equilibrium solutions for various parameter values. Solid blue curves indicate stable equilibria, and dotted red curves unstable ones (refer to §4.2.3). Fig. 2a shows the equilibrium polymer fraction *θ*\* vs. mixing parameter *χ* for a cell-free gel. (Fixed values: *N* = 100, *n*\* = 1, *τ*_0_ = 0.25, *λ* = 1.) Fig. 2b shows the effect of adding cells, with a small cell traction parameter *τ*_0_ = 0.25. Only stable equilibria (solid blue line) are present in this case. (Fixed values: *N* = 100, *n*\* = 1, *τ*_0_ = 0.25, *λ* = 1.) Fig. 2c when the cell traction is larger (*τ*_0_ = 1) the steady state polymer fraction *θ*\* increases for a given value of *χ*. (Fixed values: *N* = 100, *n*\* = 1, *τ*_0_ = 1, *λ* = 1.) Fig. 2d shows the relationship between *τ*_0_ and *χ* with the polymer equilibrium fixed at *θ*\* = 0.5. (Fixed values: *θ*\* = 0.5, *N* = 100, *n*\* = 1, *λ* = 1.) Fig. 2e illustrates how the equilibrium polymer fraction *θ*\* depends on the cell traction parameter *τ*_0_. Note that the solid green line is *τ*_0_ = 0, so equilibria to the left of this have *τ*_0_ < 0 and are not relevant. (Fixed values: *χ* = 0.75, *N* = 100, *n*\* = 1, *λ* = 1.)

With cells introduced into the system, *θ*\*increases monotonically as *χ* increases, as seen in Figs. 2b and 2c where *τ*_0_ = 0.25 and *τ*_0_ = 1 respectively. In these examples, we see that there are no longer any unstable equilibria, and that stable equilibria now exist over the spectrum of *χ* values. We see that as the traction parameter increases from *τ*_0_ = 0.25 (Fig. 2b) to *τ*_0_ = 1 (Fig. 2c), the equilibrium polymer fraction is greater for the same values of *χ*. For example, in Fig. 2b where *τ*_0_ = 0.25, at *χ* = 0 we have *θ*\* = 0.2, while in Fig. 2c where *τ*_0_ = 1, at *χ* = 0 we have *θ*\* = 0.58. Similarly, we see in these figures that with *τ*_0_ increasing from *τ*_0_ = 0.25 to *τ*_0_ = 1, the same particular equilibrium value *θ*\* is found with a decreasing value of the mixing parameter *χ*.

This relationship between *τ*_0_ and *χ* is reinforced in Fig. 2d, where *χ* is plotted against *τ*_0_ for fixed *θ*\* = 0.5. We see that *χ* decreases linearly with increasing *τ*_0_; as the cells exert more force, lower values of the interaction parameter are needed to keep the system at the same equilibrium value of polymer. Similarly, with a larger value of *χ*, less cell traction is necessary to maintain this equilibrium. This linear relationship between *χ* and *τ*_0_ when the polymer fraction is fixed can be clearly seen in equation (4.24).

In Fig. 2e it is shown that, as would be expected, larger values of *τ*_0_ correspond to larger *θ*\*, *i.e*. greater compaction in the polymer network. We note that we must have *τ*_0_ ≥ 0; therefore, the equilibria that cross the vertical line at *τ*_0_ = 0 are not biologically relevant. There is a very small branch of permissible unstable equilibria in the region approaching *θ*\* = 0, but the vast majority of initial conditions here will reach a stable steady state. In the small region where two steady states exist, gels with an initial polymer fraction above the unstable values of *θ*\* will contract to the stable equilibrium, whereas those below the unstable values will swell until the gel dissolves.

## 5 Numerical simulations

We now perform numerical simulations to investigate the behaviours predicted by our model. We consider a range of initial conditions – both uniform and non-uniform – and parameter values to better understand the emergent behaviours that can arise as a result of the interacting factors in the system. The one-dimensional governing equations (3.21a) - (3.24) are simulated in MATLAB, using finite difference methods. Central differencing is used in the velocity equation (3.21e), excluding at *X* = 1, where a one-sided difference is used for derivatives of *θ_p_*, and the boundary condition (3.23a) is used to provide a ghost point for *v_p_*. The Crank-Nicolson method is used for equations (3.21c) and (3.21d), with one-sided differences used for derivatives of *v_p_* at *X* = 1. We verified the code by comparing the simulation results with the short-time and steady state solutions derived in §4, and by checking the scheme conserves mass effectively over time. We calculated the percentage change in the mass of polymer and cells at the initial and final points in time for each of the simulations presented in this section: with a time step of *dT* = 0.0005 and spatial step of dX = 0.002, the worst-case change in mass for *θ_p_* or *n* was 0.0076%. Full details of the numerical scheme and verification checks can be found in Reoch (2020).

The aim of our simulations is to illustrate the qualitative behaviours of our model for different initial conditions and in different parameter regimes. Lacking experimental data to fit model parameters, we take the majority of parameters to be 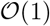. Certain initial conditions and parameters are kept fixed (see Table 1). For example, in nondimensionalising the length scale is set such that the initial length *L*(0) = 1. Similarly, when cells are present, we choose the average initial cell density as the characteristic value, so that *n_i_* = 1 for an initially uniform cell distribution. Given that 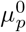 and 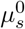 do not appear in the final set of model equations due to cancellation of terms when taking 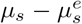 in interface condition (3.23a), we set these to zero. We set the bulk viscosities *κ_p_* and *κ_s_* to zero without loss of generality, as these terms only appear in linear combination with the dynamic viscosity parameters. The polymer chain length *N* is generally large for polymer and solution mixtures, therefore we set *N* = 100 (Rubinstein *et al*., 2003), and we set the contact inhibition parameter *λ* =1. The parameters which are varied between simulations are listed in Table 2. We note that the mixing parameter *χ* is the only term appearing in the final system of equations for which negative values are physically meaningful.

**Table 1:** Dimensionless initial conditions and parameter values which we do not change between simulations.

| Term | Symbol | Value used |
| --- | --- | --- |
| Initial length | $L_i$ | 1 |
| Contact inhibition parameter | $\lambda$ | 1 |
| Polymer chain length | $N$ | 100 |
| Polymer standard free energy | $\mu_p^0$ | 0 |
| Solvent standard free energy | $\mu_s^0$ | 0 |
| Polymer bulk viscosity | $\kappa_p$ | 0 |
| Solvent bulk viscosity | $\kappa_s$ | 0 |

**Table 2:** Dimensionless initial condition and parameter values which we may change between simulations.

| Term | Symbol | Range of values used |
| --- | --- | --- |
| Initial polymer fraction | $\theta_i$ | 0.2 - 0.7 |
| Initial cell density | $n_i$ | 0 - 1 |
| Cell traction coefficient | $\tau_0$ | 0.1 - 1 |
| Cell diffusion coefficient | $D$ | 0 - 1 |
| Mixing parameter | $\chi$ | -0.1 - 1.5 |
| Interface resistance | $\mathcal{R}$ | 0.1 - 5 |
| Drag coefficient | $\xi$ | 0 - 5 |
| Solvent dynamic viscosity | $\eta_s$ | 0.1 - 5 |

### 5.1 Cell-free gel, uniform initial conditions

In a cell-free gel (where *n* = 0), swelling or contraction is driven by the free energy of the system; gradients in chemical potentials on either side of the gel-solvent interface induce the movement of solvent and polymer. This is similar to the results presented in Keener *et al*. (2011b). In Fig. 3a, where *θ_i_* = 0.6 and *χ* = 0.75, the balance in chemical potentials *μ_p_* and *μ_s_* produces an osmotic pressure gradient, causing solvent to enter the gel from the surrounding solvent region Ω_*s*_; the gel thus swells until an equilibrium is reached with *θ*\* = 0.45 and *L*\* = 1.34. Conversely, in Figs. 3b and 3c, we see the gel contract to an equilibrium state. The free energy in the system has been altered in two different ways here to induce contraction. In Fig. 3b, we have taken the same initial conditions as Fig. 3a, but with an increased strength of mixing parameter *χ* = 1.5.

**Figure 3:**
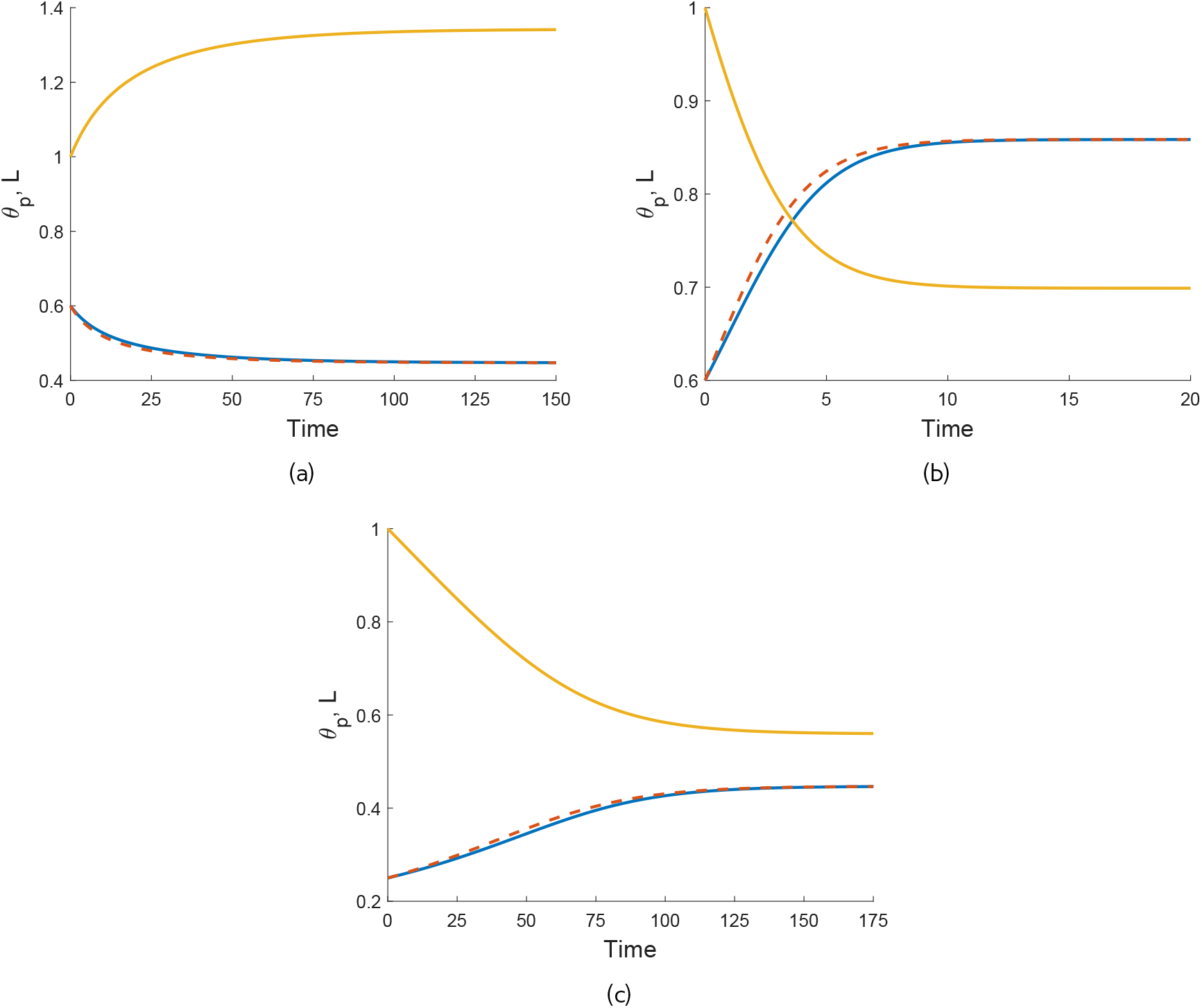
Illustrations of qualitatively different behaviours of cell-free gels. In each plot, *L*(*T*) is the solid gold line, *θ_p_*(*X* = 0) is the solid blue line, and *θ_p_* (*X* = 1) is the dashed red line. The following parameters are the same in each simulation: *n_i_* = 0, *η_s_* = 0.25, *ξ* = 0.5, 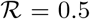. Fig. 3a: gel swells to equilibrium due to osmotic pressure when *θ_i_* = 0.6, *χ* = 0.75. Steady state: (*θ*\*, *L*\*) = (0.45, 1.34). Fig. 3b: increasing *χ*, the gel now contracts to a smaller steady state length (*θ_i_* = 0.6, *χ* = 1.5,). Steady state: (*θ*\*, *L*\*) = (0.86, 0.7). Fig. 3c: decreasing the initial polymer fraction compared to Fig. 3a changes the free energy balance such that the gel now contracts (*θ_i_* = 0.25, *χ* = 0.75). Steady state: (*θ*\*, *L*\*) = (0.45, 0.56).

In Fig. 3c, the initial fraction of polymer has been decreased to *θ_i_* = 0.25, with the value of *χ* remaining at *χ* = 0.75. The effect in both instances is to increase the initial free energy in the gel, resulting in a situation where the gradient in chemical potentials will induce solvent to flow out from the gel to balance the potentials, and hence result in a smaller equilibrium length. The gel equilibrates with *θ*\* = 0.86 and *L*\* = 0.7 in Fig. 3b, and *θ*\* = 0.45 and *L*\* = 0.56 in Fig. 3c. These equilibria all clearly satisfy the mass conservation relation *L*\**θ*\* = *θ_i_* (as indeed will all steady states found).

We note that the simulations in Figs. 3a and 3c reach the same equilibrium value, *θ*\* = 0.45, for the two different initial conditions; this corresponds to the equilibrium predicted in Fig. 2a with *χ* = 0.75 and the same fixed set of parameter values otherwise. Fig. 3b meanwhile confirms that, for the same initial conditions and parameter set, increasing the value of *χ* will result in an equilibrium with a larger polymer fraction (*θ*\* = 0.86). This is also in agreement with Fig. 2a.

We also note that the polymer fraction at *X* = 1 (shown by red dashed lines) evolves slightly faster than that at *X* = 0 (blue solid lines). This lag reflects the time taken for the solvent to flow into or away from the centre of the gel. We discuss the parameters affecting this lag and the spatial profiles of the polymer as the gel evolves in Section 5.3.

We have shown here that, in the absence of cells, the gel will swell or contract depending on the balance between chemical potential gradients across the gel-solvent interface. These behaviours echo those found by Keener *et al*. (2011b), which is expected since our gel model builds on their work. Comparing our model to that of Keener *et al*., we note that while the inclusion of boundary resistance in our model only affects the rate of the gel’s elongation or shrinking, the absence of any contribution from the standard free energy parameters 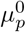 and 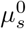 will change the final gel length and polymer fraction found here for the same set of parameters used in Keener *et al*.. We next introduce cells into the simulations to study their effect on the gel’s behaviour.

### 5.2 Cell-gel system

In Fig. 4a, we use the same gel parameters as for Fig. 3a, and introduce a cell population with weak traction (*n_i_* = 1, *τ*_0_ = 0.1; note also *D* = 0.01). We see that the gel still swells to a steady state with the cell traction parameter set at this low level. However, compared to the simulation in Fig. 3a, the final size of the gel is now smaller (equilibrating here at *θ*\* = 0.54, *n*\* = 0.91, *L*\* = 1.1), indicating that the cells are exerting some contractile force that counters the expansion due to osmotic effects. We increase the traction parameter to *τ*_0_ = 1 in Fig. 4b; once cell traction is increased over a certain threshold, the gel will switch from expansion to contraction. In this instance, the cell traction stresses are stronger than the chemical potential gradient, and as the cells compact the polymer network, solvent is squeezed from the gel until it reaches a steady state once the mechanical forces are in balance, where *θ*\* = 0.86, *n*\* = 1.44, *L*\* = 0.69. We have therefore established that introducing cells into a gel that would otherwise swell can induce a switch in behaviour, resulting in a significantly different outcome for the gel. As in Section 5.1, the equilibria found here satisfy the mass conservation relations *L*\**θ*\* = *θ_i_*, *L*\**n*\* = 1.

**Figure 4:**
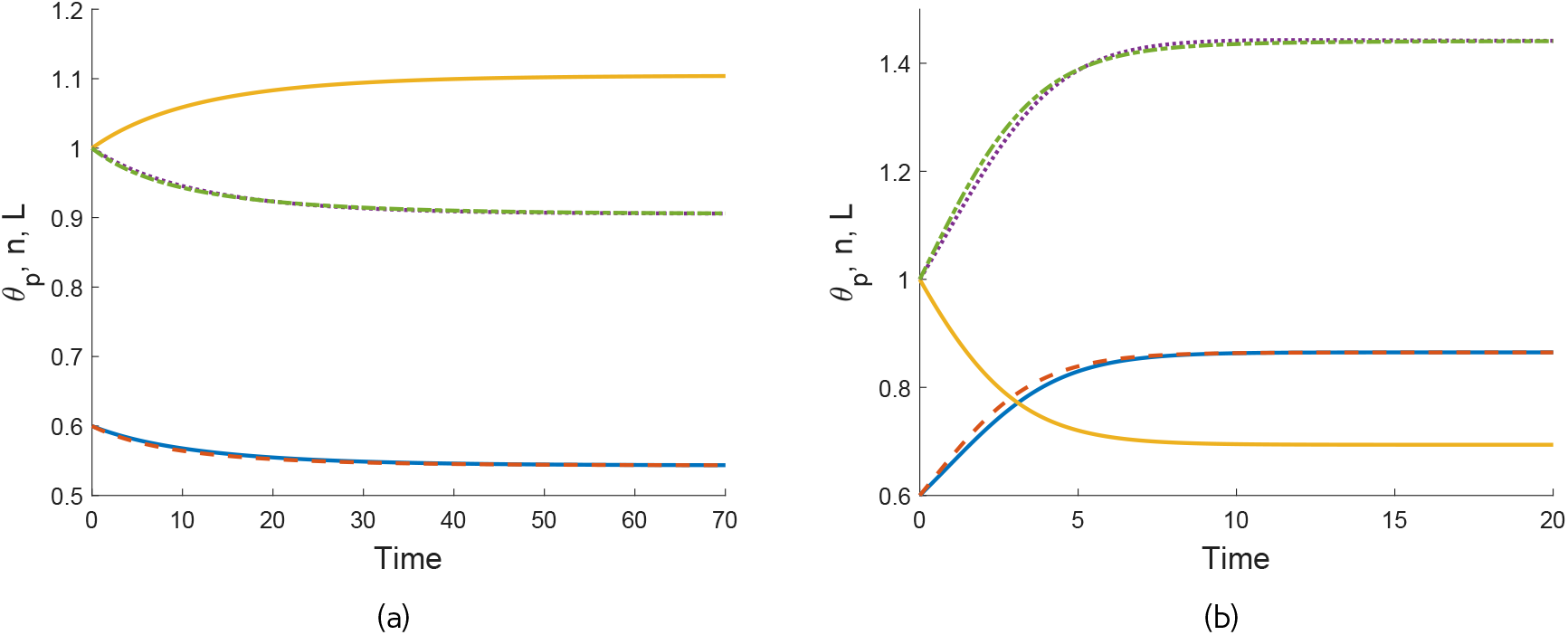
Time evolution of a cell-gel system. Colour key: *L*(*T*) - gold; *θ_p_*(*X* = 0) - blue, *θ_p_*(*X* = 1) - dashed red; *n*(*X* = 0) - dotted purple; *n*(*X* = 1) - dash-dotted green. Fig. 4a: when the cell traction is relatively weak, the gel still swells to an equilibrium state. (Parameter values: *θ_i_* = 0.6, *n_i_* = 1, *χ* = 0.75, *η_s_* = 0.25, *ξ* = 0.5, 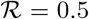, *τ*_0_ = 0.1, *D* = 0.01.) (*θ*\*, *n*\*, *L*\*) = (0.54, 0.91, 1.1). Fig. 4b: With increased cell traction, the gel switches to contraction as the cell-induced forces are stronger than the osmotic pressure. (Parameter values: *θ_i_* = 0.6, *n_i_* = 1, *χ* = 0.75, *η_s_* = 0.25, *ξ* = 0.5, 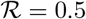, *τ*_0_ = 1, *D* = 0.01.) (*θ*\*, *n*\*, *L*\*) = (0.86, 1.44, 0.69).

The time taken to equilibrate is noticeably different across the simulations seen so far, as the rate at which the gel evolves is affected by the strength of a number of competing forces. For example, we see in Fig. 3a that equilibrium is reached at approximately *T* = 130, while in Fig. 3b, due to the larger value of *χ*, not only does the gel contract, but it equilibrates by *T* = 15. In a case like that presented in Fig. 3b, where the free energy alone induces gel contraction, adding cells to this gel will lead to a steady state being reached more quickly (result not shown). Therefore, the magnitude of parameters like the interaction energy *χ* and cell traction *τ*_0_ will affect the time taken to reach a steady state. Alongside this, mechanical factors like drag and viscosity will impact the gel’s temporal evolution.

### 5.3 Effects of mechanical parameters and diffusion on gel evolution

We now study how the ratios of drag *ξ*, resistance 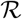, and solvent viscosity *η_s_* relative to polymer viscosity *η_p_* affect the rate at which a gel evolves to equilibrium and the manner in which it does so spatially. In the simulations presented in Figs. 5a - 6b, we take a gel with the same initial conditions, free energy parameters, and cell force parameters as that presented in Fig. 4b; this gel will therefore reach the same equilibrium regardless of parameters like drag and viscosity (given that the equilibrium is determined by equation (4.5), in which these mechanical parameters do not appear). We first set *ξ*, 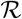 and *η_s_* to be small, indicating that these effects are insignificant relative to polymer dynamic viscosity, and as such, polymer dynamic viscosity is the dominant mechanical characteristic; we will refer to this as the base case in the comparisons that follow. We note that due to the scaling used here, we have effectively set *η_p_* = 1. To better understand the impact of each parameter on the gel’s evolution, we then change one of *ξ*, 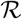 and *η_s_* in turn while holding the others constant. We take *D* = 0 here so that diffusion has no impact on the cell and polymer distributions. The gel reaches the same steady state in each case (*θ*\* = 0.86, *n*\* = 1.44, *L*\* = 0.69), albeit at different times and with different lags between *X* = 1 and *X* = 0.

**Figure 5:**
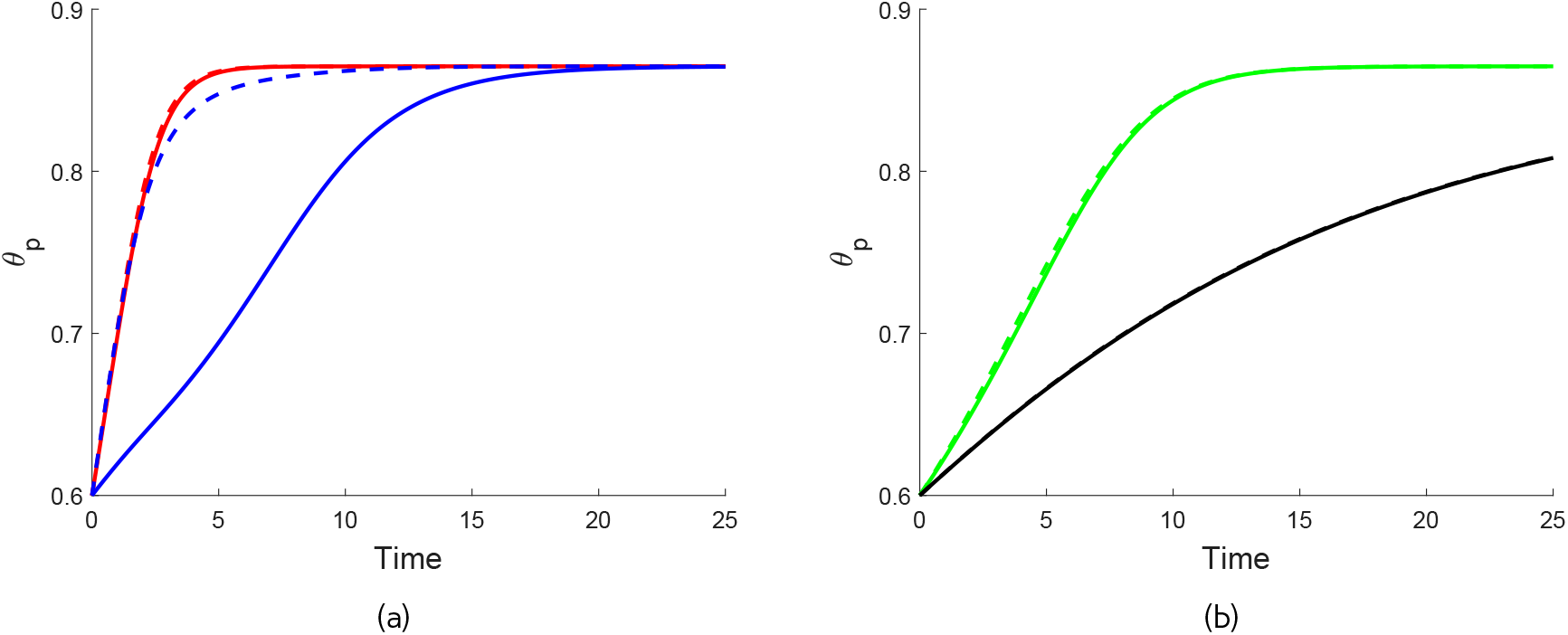
Effects of drag, resistance and solvent viscosity on gel behaviour (with *θ_i_* = 0.6, *n_i_* = 1, *χ* = 0.75, *η_s_* = 0.1, *τ*_0_ = 1, *D* = 0 fixed). The base case 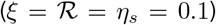 is shown by the red solid *θ_p_*(*X* = 0) and *θ_p_*(*X* = 1) dashed red curves, respectively, in Fig. 5a. The corresponding spatial profiles for *θ_p_* at *T* = 0, 0.2, 0.5, 1, 1.5, 2, 3, 5 are shown in Fig. 6a. For large drag (*ξ* = 5, 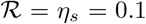), the change in the polymer fraction *θ_p_* is much slower at *X* = 0 compared to *X* = 1 (blue solid and dashed curves, respectively, in Fig. 5a) as the solution encounters greater resistance as it flows through the gel to the interface. For large resistance (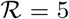, *ξ* = *η_s_* = 0.1) - see green curves in Fig. 5b - the rates of change in the polymer fraction are very similar in the centre and at the edge of the gel, but both are slower than the base case due to the additional resistance to outflow of solution at the interface. A similar effect is seen for large solvent viscosity (*η_s_* = 5, 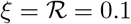) - black curves in Fig. 5b.

**Figure 6:**
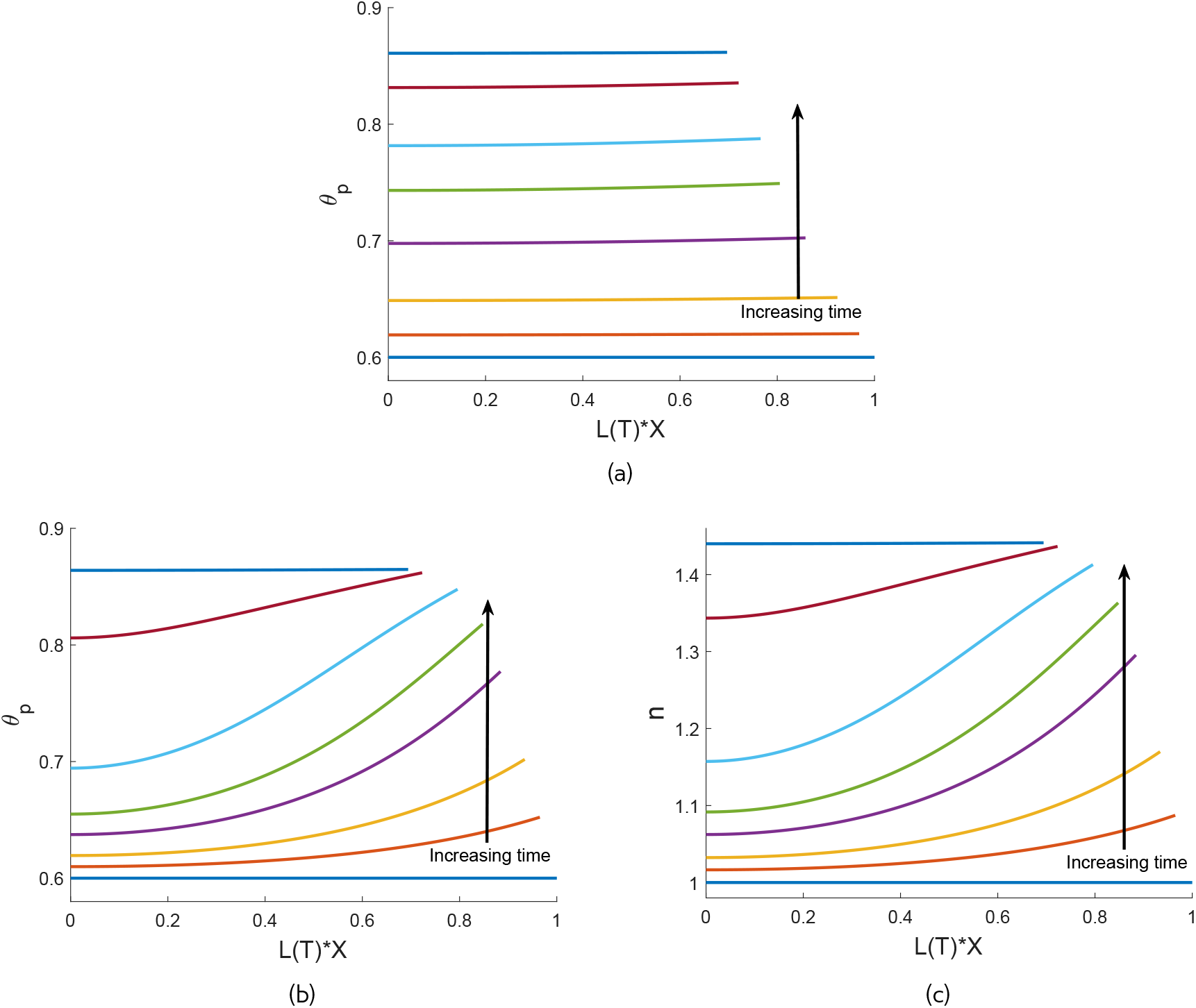
Evolution of the polymer and cell distributions for the base and large-drag cases from Fig. 5a. Fig 6a shows the profile for *θ_p_* at *T* = 0, 0.2, 0.5, 1, 1.5, 2, 3, 5 for the base case parameter values. Figs. 6b and 6c show the spatial profiles for *θ_p_* and *n*, respectively, for the high-drag case.

In Fig. 5a, we compare the evolution of *θ_p_* for the base case with 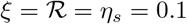 (shown by the red solid line for *X* = 0 and the red dashed line for *X* = 1) and for a gel with large drag, where *ξ* = 5 and 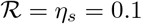 (shown by the blue solid line for *X* = 0 and blue dashed line for *X* = 1). We see that increasing the drag coefficient slows down the evolution of the polymer fraction (with the gel equilibrating at *T* ≈ 25 with large *ξ* compared to *T* ≈ 10 with small *ξ*). Furthermore, for large drag, *θ_p_* changes at a much slower rate at *X* = 0 than at *X* = 1, while there is little difference in *θ_p_* between *X* = 0 and *X* = 1 when drag is small. This is reflected in Figs. 6a and 6b, which show the spatial profiles for *θ_p_* at increasing points in time for the base case and large drag case respectively. With polymer viscosity dominant in Fig. 6a, the gel evolves across the spatial domain in a largely uniform manner. In Fig. 6b, much stronger spatial variations are evident, reflecting that, while the gel evolves quickly at the interface, it takes much longer for solvent to flow through the domain due to the extra resistance when drag is large.

In Fig. 5b we similarly compare the gel’s behaviour with interface resistance 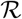 and solvent viscosity *η_s_* each large relative to polymer viscosity. For large resistance (shown by the green solid and dotted curves for *X* = 0 and *X* = 1 respectively), we take 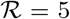, *ξ* = *η_s_* = 0.1, while for large viscosity (shown by the black solid and dotted curves for *X* = 0 and *X* = 1 respectively), we set *η_s_* = 5, 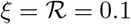. Note that this figure should be compared to the base case given by the red curves in Fig. 5a. Increasing the resistance parameter 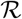, the rate of change of *θ_p_* is slowed at *X* = 1 compared to the base case (with equilibrium now reached at *T* ≈ 20). This reflects the fact that the boundary of the gel is less permeable to fluid flow with larger 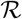. In this case, the polymer fraction remains almost uniform across the spatial domain, *i.e*. there is no additional lag induced between *X* = 1 and *X* = 0 with large 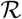. Large solvent viscosity *η_s_* has the effect of slowing down gel contraction further still, with the gel not equilibrating until *T* ≈ 80 (note that equilibrium for the large *η_s_* case is not shown in Fig. 5b). As with 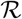, in this case there is minimal lag between the evolution at *X* = 1 and *X* = 0.

We now study the effect of non-zero diffusion on the gel’s behaviour. As diffusion increases, we expect to see more uniform spatial profiles in the cell density as well as the polymer fraction, as cells spread more evenly across the gel through random motion. In Fig. 6b we showed the spatial profiles of *θ_p_* for a gel with a large drag coefficient and zero diffusion. In Fig. 6c, we see that the spatial distribution of cells in this gel is similarly non-uniform over much of the gel’s evolution. We now take this same gel with large drag, but introduce cell diffusion, setting *D* = 0.005; the time evolution in this case is shown in Fig. 7a. In Fig. 7b, we see that with large drag, the cell density initially increases rapidly at *X* = 1 (like in Fig. 6c). Over time, cells move down their density gradient towards *X* = 0 due to the diffusion term now present. The cells become more dense in this region, eventually overshooting the equilibrium value (see *n* in Fig. 7a); however, as time progresses, the diffusive movement leads *n* to converge to its equilibrium value across the domain. Increasing diffusion further to *D* = 1 in Fig. 7c, the strength of the random motion is such that the cells remain well spread spatially for all time. We note that in this case, the polymer still evolves with a lag across the spatial domain, like seen in Fig. 6b (result not shown).

**Figure 7:**
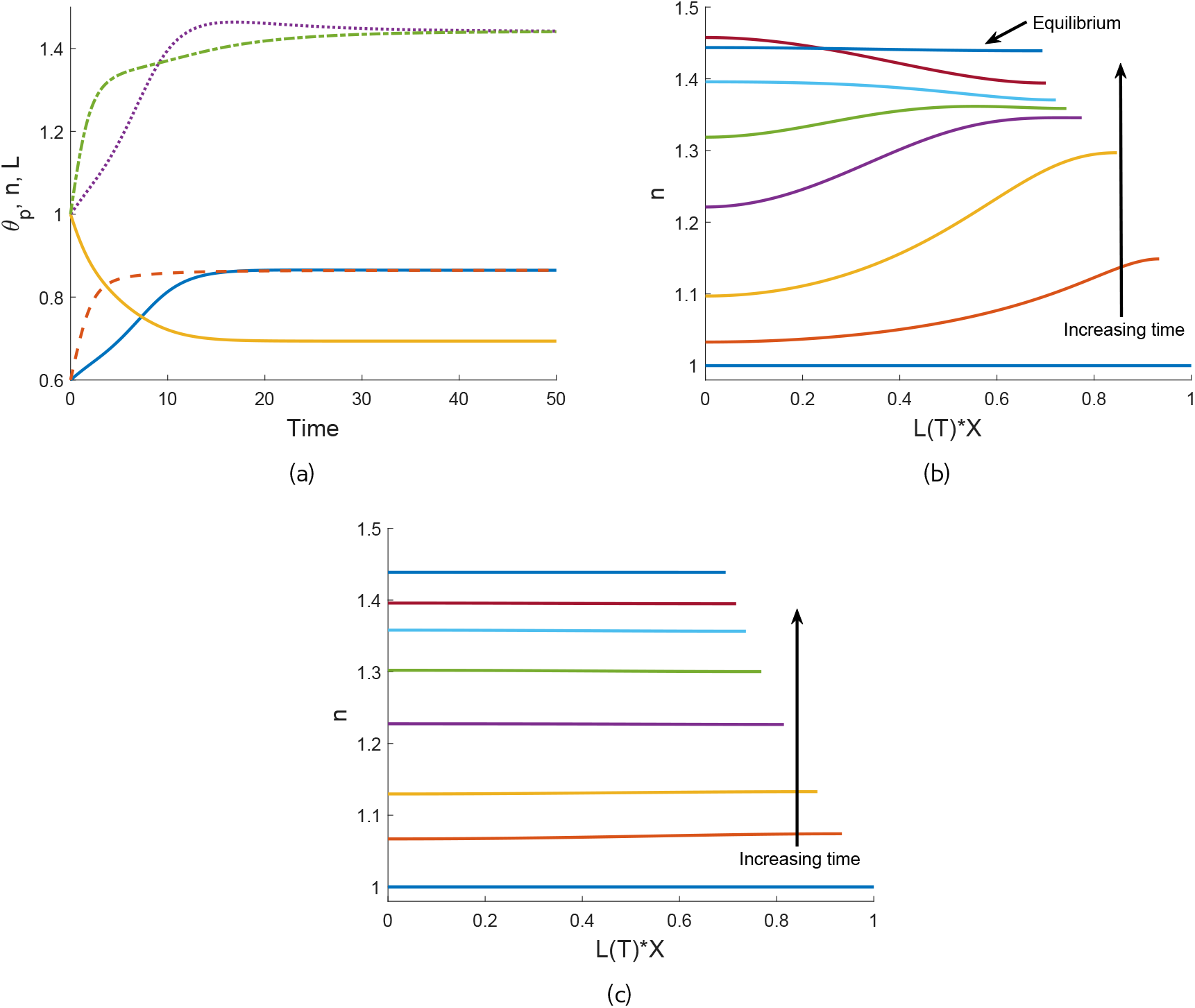
The effect of cell diffusion on gel behaviour. Fig. 7a - with small diffusion, the cell density and polymer fraction increase slightly above their equilibrium values at *X* = 0 before slowly reducing to the steady state. Colour key: *L*(*T*) - gold; *θ_p_*(*X* = 0) - blue; *θ_p_*(*X* = 1) - dashed red; *n*(*X* = 0) - dotted purple; *n*(*X* = 1) - dash-dotted green. Parameter values: *ξ* = 5 and *D* = 0.005 with other parameters as in Fig. 5a. 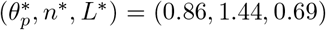. Fig. 7b shows the spatial profiles for *n* for the solutions shown in Fig. 7a at times *T* = 0, 1, 3, 6, 8, 10, 14, 40. Fig. 7c shows the effect of increasing the cell diffusion to *D* = 1 (other parameter values as in Fig. 7a). Comparing Figs. 7b and 7b, we see the cells now remain almost uniformly distributed as the gel contracts. Profiles are plotted (from bottom to top) at *T* = 0, 1, 2, 4, 6, 8, 10, 18.

### 5.4 Reduced initial polymer fraction

Taking a smaller initial polymer fraction, we can see different dynamics emerge in the evolution of the gel. Setting *θ_i_* = 0.2 and all other parameters as in Fig. 4b, we see in Figs. 8a and 8b that the polymer fraction and cell density evolve slowly over the beginning phases (*T* = 0 to *T* ≈ 2), before a period of rapid increase (*T* ≈ 2 to *T* ≈ 7), which slows down again as the gel moves towards its steady state (T ≈ 7 to *T* ≈ 12). From this lower initial value of *θ_p_*, much greater contraction is evident in the gel, which reaches the steady state *θ*\* = 0.91, *n*\* = 4.54, *L*\* = 0.22. The evolution of the gel length here is quite rapid until the gel approaches its steady state length (*T* ≈ 7). With a larger initial polymer fraction, *θ_i_* = 0.4, the gel reaches the steady state (*θ*\*, *n*\*, *L*\*) = (0.89, 2.23, 0.45) (result not shown). In Fig. 4b, we saw that with *θ_i_* = 0.6, the resulting steady state was (*θ*\*, *n*\*, *L*\*) = (0.86, 1.44, 0.69). This demonstrates that, with the initial cell density constant, there is a negative correlation between the initial fraction of polymer in the gel and the degree to which both the gel contracts (seen in *L*\*) and the polymer compacts (seen in *θ*\*). The reason is that for spatially uniform steady states and initial conditions, by mass conservation we must have *L*\**θ*\* = *θ_i_*, *L*\**n*\* = 1; this implies that *n*\* = *θ*\*/*θ_i_*. We see that the decrease in *θ*\* and *n*\* with *θ_i_* increasing, as in the examples above, requires an increase in the equilibrium length *L*\* to conserve mass for these quantities.This negative correlation was also observed in experiments presented in Stevenson *et al*. (2010). This example is discussed further in relation to other models in Section 6.

**Figure 8:**
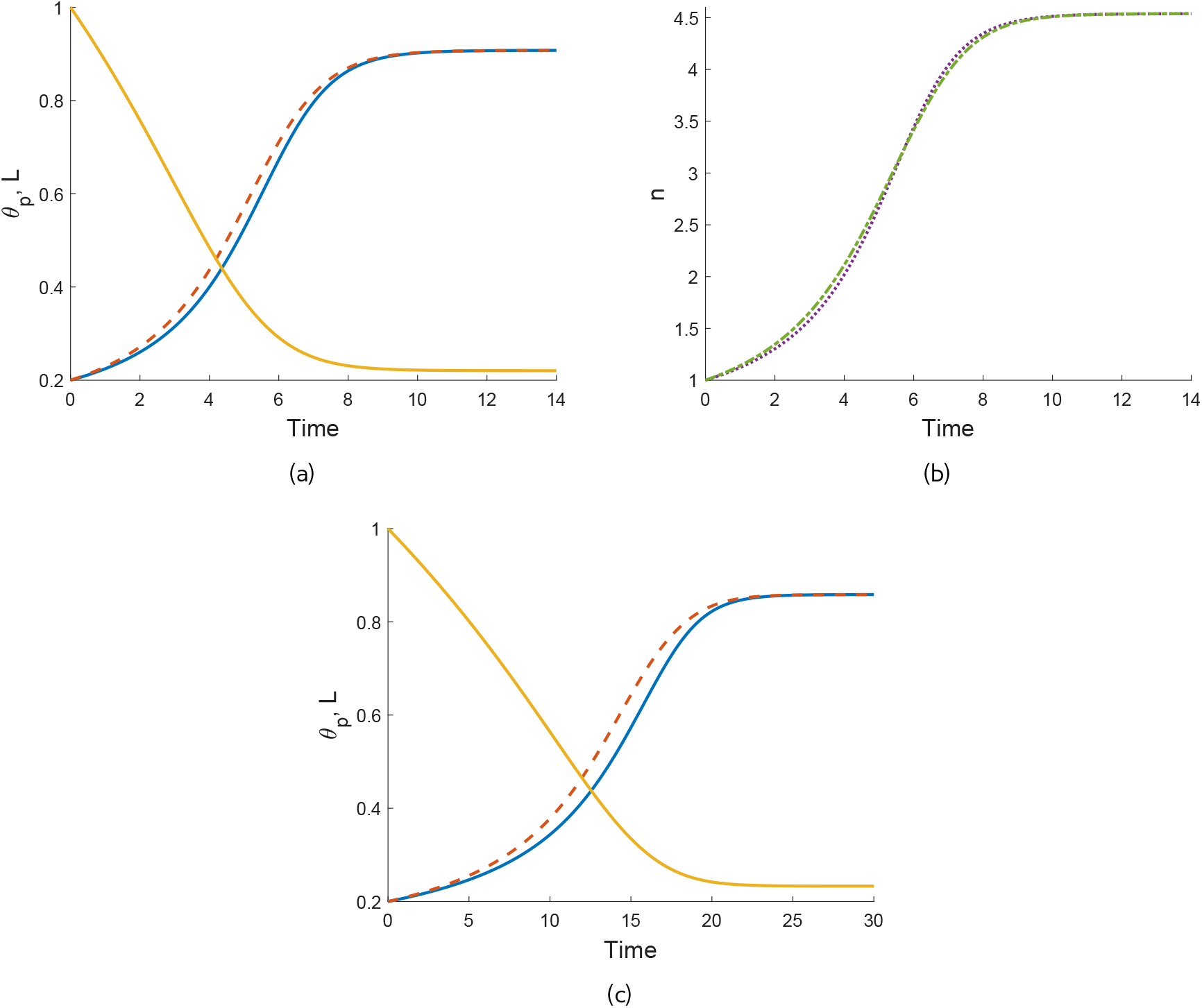
The effects of reduced initial polymer fraction on gel contraction. Fig. 8a - with a small initial polymer fraction *θ_i_* = 0.2, a significant degree of contraction occurs, resulting in a much smaller gel at equilibrium. Colour key: *L*(*T*) - gold; *θ_p_*(*X* = 0) - blue; *θ_p_*(*X* = 1) - red. Parameter values: *n_i_* = 1, *χ* = 0.75, *η_s_* = 0.25, *ξ* = 0.5, 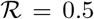, *τ*_0_ = 1, *D* = 0.01. 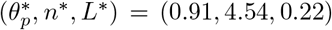. Fig. 8b shows the corresponding plots for *n*. Fig. 8c shows similar behaviour for a cell-free gel. Parameter values: *θ_i_* = 0.2, *n_i_* = 0, *χ* = 1.5, *η_s_* = 0.25, *ξ* = 0.5, 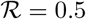. 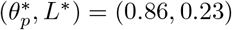.

We note that we see similar behaviour take place in a cell-free gel. For example, amending the example in Fig. 8a such that *θ_i_* = 0.2, *n_i_* = 0, *χ* = 1.5, the gel evolves in a similar manner, with a slow initial phase of polymer compaction followed by rapid evolution and then slowing near the steady state (see Fig. 8c). In this instance, the gel equilibrates with *θ*\* = 0.86, *L*\* = 0.23. Furthermore, taking *θ_i_* = 0.4 and *θ_i_* = 0.6, we reach steady states of (*θ*\*,*L*\*) = (0.86, 0.47) and (*θ*\*,*L*\*) = (0.86, 0.70) respectively (result not shown for *θ_i_* = 0.4; see Fig. 3b for *θ_i_* = 0.6). While the equilibrium fraction of polymer is the same regardless of the initial condition in the cell-free case (provided the same free energy parameters are used), we see that the change in gel length is also negatively correlated with the initial polymer fraction here. This is clear from the mass conservation condition *L*\**θ*\* = *θ_i_*, given *θ*\* is fixed by the free energy parameters in the cell-free case. These results indicate that the presence of cells is therefore necessary to see the negative correlation between initial and final polymer fractions.

### 5.5 Non-uniform initial conditions

The numerical simulations presented thus far have been performed using spatially uniform initial conditions. Despite spatial variations being evident while the gel is evolving, these initial conditions eventually produce spatially uniform steady states. This matches previous work such as Keener *et al*. (2011b) looking at cell-free models, wherein only spatially uniform equilibria are found.

We now consider examples with spatially non-uniform initial conditions, finding that these initial conditions can result in spatially varying equilibrium solutions. This is a novel behaviour arising in our model from the presence of cells. We will consider non-uniform initial conditions in both the polymer and cells separately.

We first evaluate a cell-free gel with a spatially varying initial polymer distribution; this allows us to establish a baseline against which we can evaluate the impact of cells. We take a gel as specified in Fig. 3a where *θ_i_* = 0.6 and *χ* = 0.75, for which the gel swells to an equilibrium with *θ*\* = 0.45 and *L*\* = 1.34. We add a spatially varying component to the initial condition for the polymer fraction, such that *θ_i_* = 0.6 + 0.025 cos(*πX*); this corresponds to a gel where the polymer is slightly bunched at the gel’s centre (where *X* = 0) and less than the mean value at the gel’s edge (where *X* = 1). We note that this initial condition satisfies the symmetry condition *∂θ_p_*/*∂X* = 0 at *X* = 0. In Fig. 9a, we see that this gel swells to the same steady state as for uniform *θ_i_* = 0.6, and evolves on a similar time scale (see Fig. 3a for comparison). In Fig. 9b, we see the spatial distribution of polymer across the length of the gel at increasing points in time; the polymer, initially more concentrated towards *X* = 0, smooths out over time as the gel swells, eventually becoming uniform as it expands to its steady state where *θ*\* is constant. Therefore, in the simulations we have seen, spatial variations in the initial polymer distribution in a cell-free gel do not affect the equilibrium outcome.

**Figure 9:**
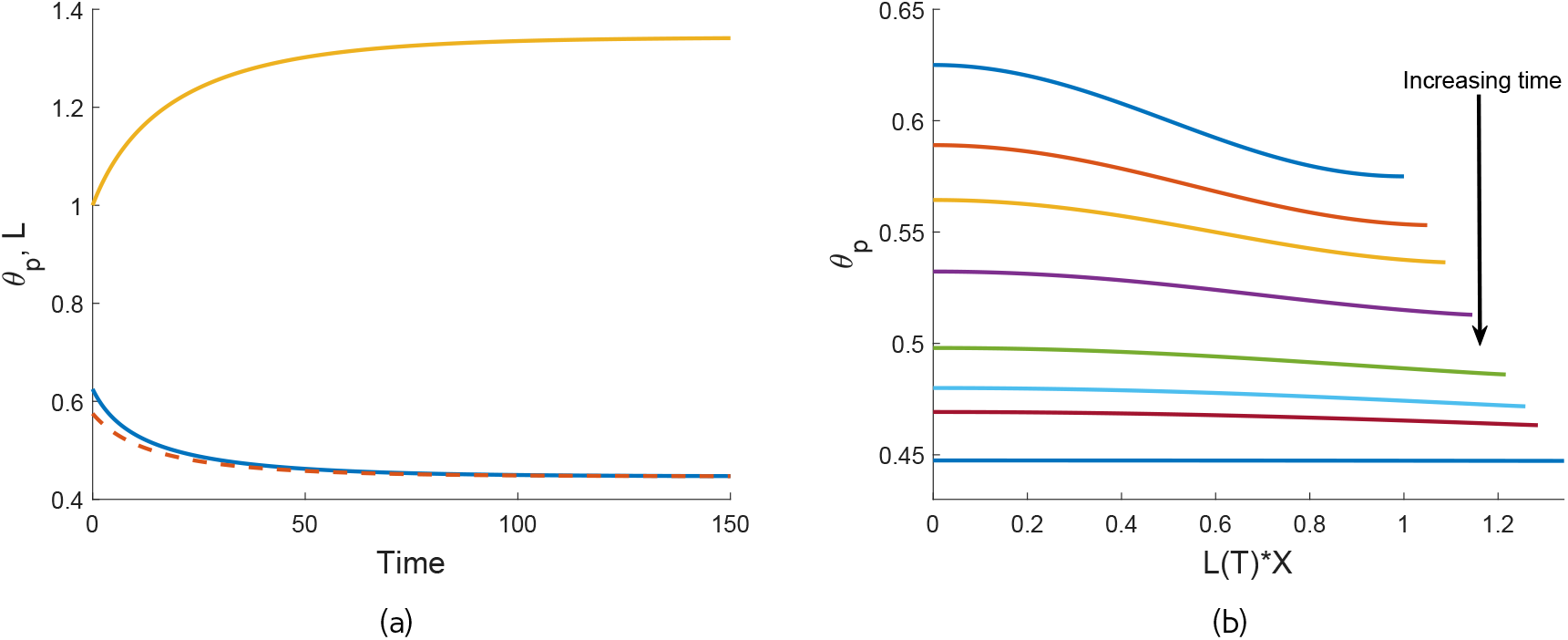
Behaviour of a cell-free gel with non-uniform initial polymer fraction which swells to a spatially uniform steady state. Parameter values: *θ_i_* = 0.6 + 0.025 cos(*πX*), *n_i_* =0, *χ* = 0.75, *η_s_* = 0.25, *ξ* = 0.5, 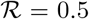. 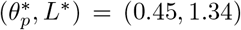. Fig. 9a shows the evolution in time (colour key: *L*(*T*) - gold; *θ_p_*(*X* = 0) - blue; *θ_p_*(*X* = 1) - dashed red), whilst Fig. 9b shows the corresponding spatial profiles for *θ_p_* at *T* = 0, 2.5, 5, 10, 20, 30, 40, 150.

We now take the same gel with a spatially varying polymer initial condition and include a cell population where *n_i_* = 1 and *τ*_0_ = 1, noting that *D* = 0. The time evolution for this gel is shown in Fig. 10a; we see that the gel, which swelled in the absence of cells due to osmotic effects, now contracts to an equilibrium with spatially non-uniform solutions for both the polymer fraction and cell density. The mean equilibrium values of *θ_p_* and *n* here, 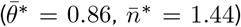, are the same as the steady state found in Fig. 4b. Figs. 10b and 10c show the spatial distributions of *θ_p_* and *n* respectively over time. In contrast to the cell-free case, the cell forces present in the system are stronger than the chemical potentials, and so induce the gel to contract. We see that the polymer is initially less compacted towards *X* = 1; it must therefore contract more in this region to move to its steady state value. As the fraction of polymer increases in this region, cells then become more concentrated, which reinforces this non-uniform evolution by pulling more polymer and cells towards the edge of the gel. The presence of non-zero drag also contributes to the formation of the spatial gradients seen, as discussed earlier. With *D* = 0, there is no requirement for the cells to even out over time, and therefore, we see a non-uniform distribution remaining at equilibrium. By the end of the process, the polymer fraction has ended slightly larger at *X* = 1 than at *X* = 0, reflecting the greater density of cells in that region.

**Figure 10:**
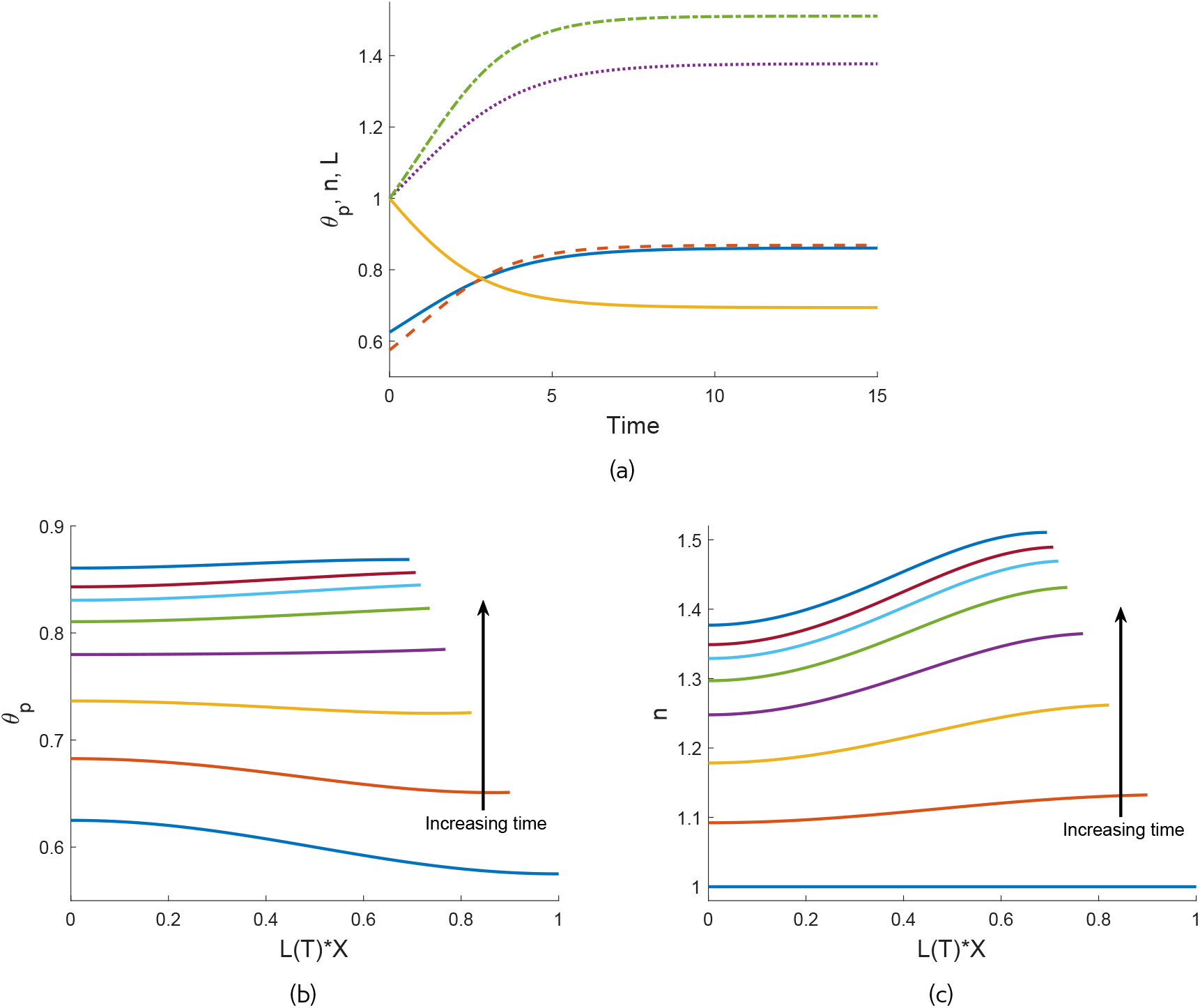
Evolution of a cell-gel system with non-uniform initial polymer fraction, where the gel contracts to a spatially-varying steady state. Parameter values: *θ_i_* = 0.6 + 0.025cos(*πX*), *n_i_* = 1, *χ* = 0.75, *η_s_* = 0.25, *ξ* = 0.5, 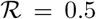, *τ*_0_ = 1, *D* = 0. 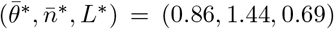. Fig. 10a shows the time evolution (colour key: *L*(*T*) - gold; *θ_p_*(*X* = 0) - blue; *θ_p_*(*X* = 1) - dashed red; *n*(*X* = 0) - dotted purple; *n*(*X* = 1) - dash-dotted green) whilst Figs. 10b and 10c show the corresponding spatial profiles for *θ_p_* and *n*, respectively, at *T* = 0, 1, 2, 3, 4, 5, 6, 15.

We now take the same system and change the spatial perturbation to the initial condition from the polymer fraction to the cell density, such that our initial conditions are *n_i_* = 1 + 0.05 cos(*πX*), *θ_i_* = 0.6. This corresponds to a gel where the cells are now initially more densely seeded at *X* = 0 (*i.e*. the centre of a gel that is symmetric about the origin). In the time evolution for this system, shown in Fig. 11a, we see the gel reaches a steady state with the same mean polymer fraction and cell density as the previous example, albeit with different spatial distributions. Fig. 11b confirms that the polymer velocity *v_p_* goes to zero across the spatial domain over time, demonstrating that the gel is maintaining its equilibrium state. The spatial profiles here are displayed in Figs. 11c and 11d. The presence of drag creates a slight increase in both *θ_p_* and *n* at *X* = 1 while the gel evolves. As the gel approaches its steady state, the evolution at *X* = 1 slows while the polymer fraction and cell density continue to increase across the rest of the domain. Greater cell concentrations around *X* = 0 result in polymer being pulled to the gel centre, and finally, a higher polymer fraction in this region at equilibrium. At this resulting steady state, we see that the amplitude of the cell profile is greater than the amplitude of *n_i_* (the amplitude is approximately 0.04 at equilibrium and 0.025 initially), while the polymer fraction is slightly larger at *X* = 0 compared to *X* = 1, in contrast to the previous example.

**Figure 11:**
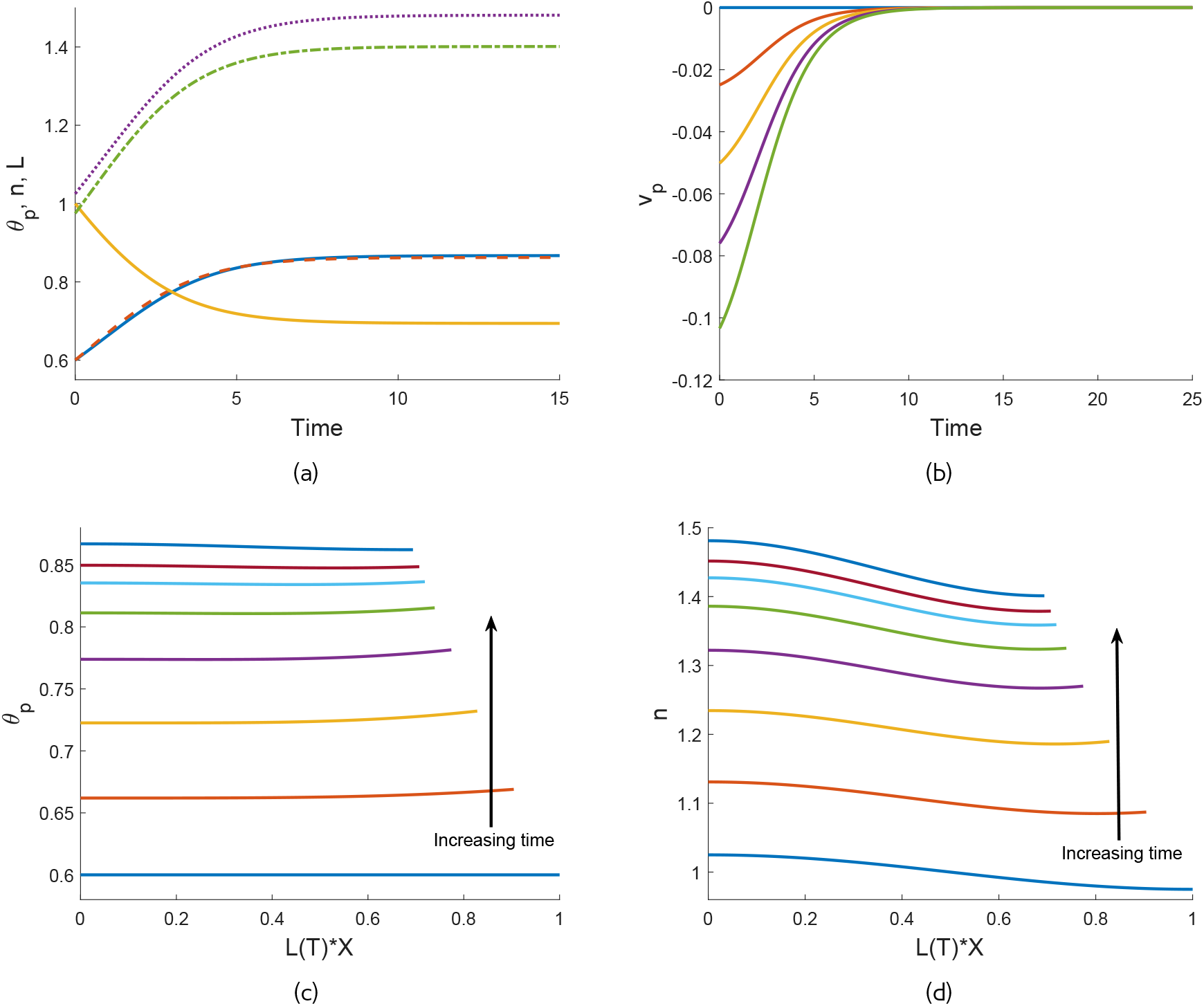
Contraction of a cell populated gel with non-uniform initial cell density and uniform initial polymer fraction. Parameter values: *θ_i_* = 0.6, *n_i_* = 1 + 0.025 cos(*πX*), *χ* = 0.75, *η_s_* = 0.25, *ξ* = 0.5, 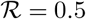, *τ*_0_ = 1, *D* = 0. 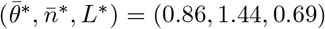. Fig. 11a shows the time evolution of *θ_p_*, *n* and *L* (colour key: *L*(*T*) - gold; *θ_p_*(*X* = 0) - blue; *θ_p_*(*X* = 1) - dashed red; n(*X* = 0) - dotted purple; *n*(*X* = 1) - dash-dotted green). Fig. 11b displays the time-evolution of the velocity at different points in the domain, showing how goes to zero across the spatial domain as the gel reaches its spatially varying equilibrium before *T* = 20. (Colour key: *v_p_*(0, *T*)- blue; *v_p_*(0.25, *T*) - red; *v_p_*(0.5, *T*) - yellow; *v_p_*(0.75, *T*) - purple; *v_p_*(1, *T*) - green.) Figs. 11c and 11d display the corresponding spatial profiles for *θ_p_* and *n*, respectively, showing that spatial variations persist to equilibrium (times plotted: *T* = 0, 1, 2, 3, 4, 5, 6, 15).

These non-uniform equilibria have been found with diffusion *D* = 0. As shown in Section 4, for the gel to equilibrate with *D* ≠ 0, *n*\* and *θ*\* must be spatially uniform. We have confirmed numerically that adding diffusion to these simulations will always result in a spatially uniform steady state on a long enough time horizon, with the additional random cell motion smoothing the cell profile, and subsequently, polymer profile as well (see (Reoch, 2020) for examples).

### 5.6 Oscillating behaviour

Our model exhibits a novel behaviour where the cell density and polymer fraction will spatially oscillate as the system evolves, *i.e*. parts of the gel will switch back and forth between swelling and contraction over time; this is displayed in Fig. 12a. In this case, uniform initial conditions are used, with *θ_i_* = 0.5 and *n_i_* = 1, while there is a negative mixing parameter *χ* = −0.1 (indicating that the polymer and solvent would prefer to mix) and fairly strong cell traction *τ*_0_ = 0.8. This behaviour emerges from the combination of parameters leading to the cell traction stress being finely balanced with the free energy. We have found that it also requires both the drag parameter *ξ* and the resistance parameter 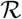 to be sufficiently large (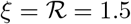 here for example), otherwise these oscillations are not evident. We note that this behaviour occurs on a very long time scale and that the gel eventually dissolves in this situation (with *θ_p_* → 0). The length of the gel increases monotonically over time (result not shown).

**Figure 12:**
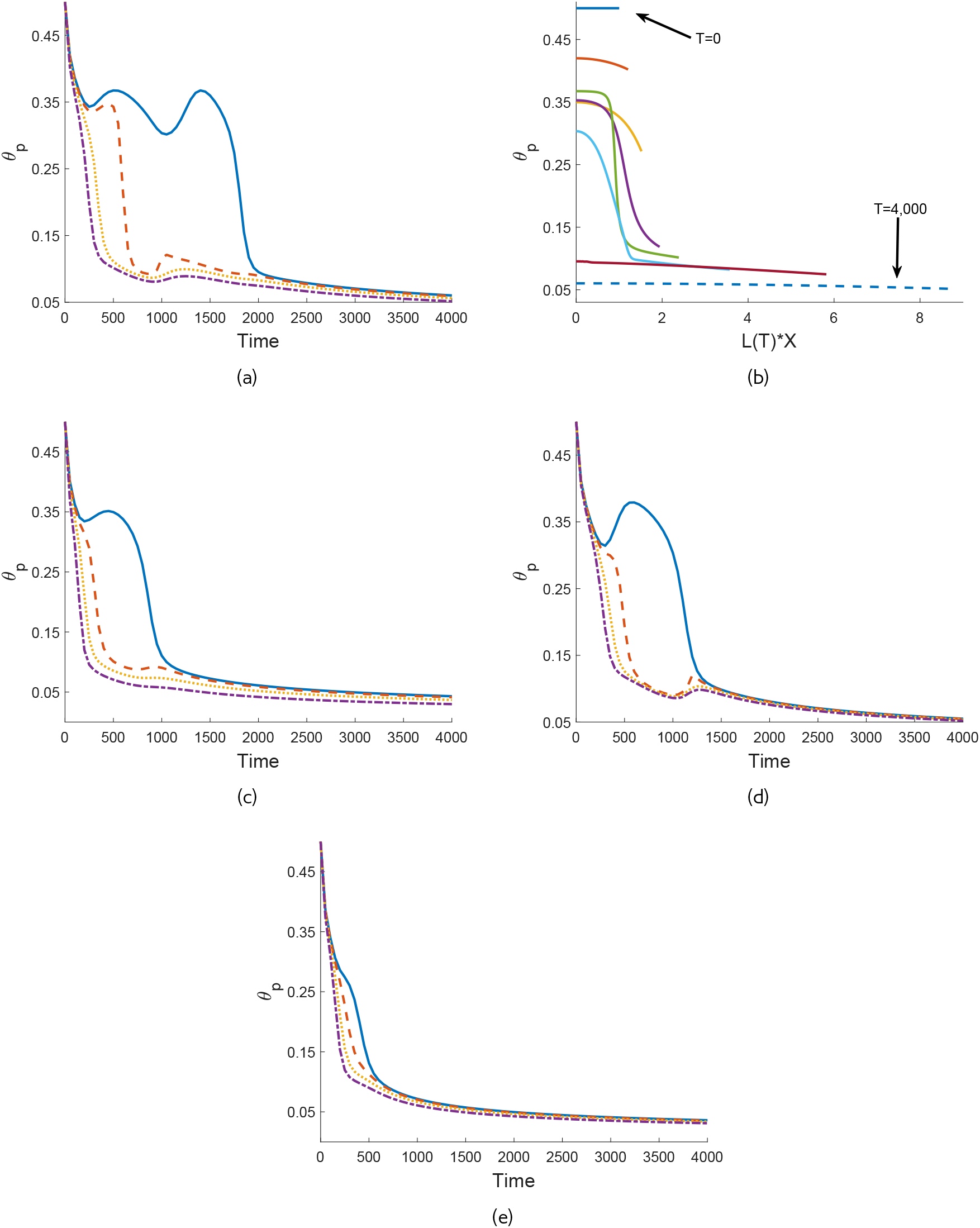
Evolution of a swelling gel with oscillating behaviour; the gel eventually dissolves as it continues to swell. Parameter values: *θ_i_* = 0.5, *n_i_* = 1, *χ* = −0.1, *η_s_* = 0.1, *ξ* = 1.5, 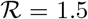, *τ*_0_ = 0.8, *D* = 0. Fig. 12a shows the time evolution of *θ_p_* (colour key: *θ_p_*(*X* = 0) - blue; *θ_p_*(*X* = 0.33) - dashed red; *θ_p_*(*X* = 0.66) - dotted gold; *θ_p_*(*X* = 1) - dash-dotted purple). Fig. 12b shows the corresponding spatial profiles at *T* = 0, 50, 200, 350, 500, 1000, 2000, 4000 (lines with increasing length correspond to increasing points at time). Reducing the resistance parameter (to 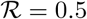) or the drag (to *ξ* = 0.5) reduces the oscillation in *θ_p_*, as shown in Figs. 12c and 12d, respectively (note colour key and other parameter values are as for Fig. 12a). Reducing both (to 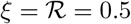) eliminates the oscillations altogether - see Fig. 12e.

This behaviour comes about as a result of the competition between osmotic effects working to expand the gel and cell traction acting to contract it. The interface resistance slows the evolution at *X* = 1, while due to the presence of drag, steep gradients develop in the polymer fraction and cell density as the gel swells, with *θ_p_* and *n* decreasing most near *X* = 1. These significant gradients can be seen in the spatial profiles for *θ_p_* shown in Fig. 12b. Large variations in the cell density are similarly evident between the two ends of the spatial domain (result not shown). The greater density of cells around *X* = 0 produces a gradient such that the cell force starts to pull polymer back towards *X* = 0. The gel then contracts locally in this region, while still swelling across the domain towards *X* = 1. Eventually, the chemical potential gradients are such across the domain that the gel starts swelling for all *X* again, with these local fluctuations repeating once more as the gel slowly expands in a non-uniform manner. Over a long enough time frame, the gel eventually dissolves.

In Figs. 12c - 12e, we see the effect of reducing the interface resistance and drag in these simulations. With resistance and drag each taken separately to be small (Figs. 12c and 12d respectively), we see reduced oscillations in the gel; however, this behaviour still occurs. When both parameters are taken to be small, the oscillations are no longer present and the gel dissolves in a more typical manner (see Fig. 12e). Similarly, with very small diffusion (*e.g*. *D* ≈ 0.0001) the oscillations occur, but larger diffusion coefficients smooth out spatial gradients in the cell density and so prevent this oscillating behaviour from occurring (results not shown).

## 6 Discussion

In this paper, we have presented a new model for cell-induced gel contraction, and studied its behaviour in a 1D Cartesian geometry. This has allowed us to develop a thorough understanding of the conditions under which the gel equilibrates, the conditions affecting the early time behaviour and the stability of the system, and, through numerical analysis, the qualitative behaviours that can occur. Throughout, we have seen that the balance between chemical potentials and cell-derived forces is crucial to determining the gel’s behaviour. We have shown that the presence of cells can cause a gel that would otherwise swell to contract; meanwhile, sufficiently strong osmotic forces can cause a gel to swell even with cells present. Moreover, the initial fraction of polymer was shown to negatively correlate with the final polymer fraction in cell-gel systems, and negatively correlate with the final gel length with or without cells present.

In Section 4, we studied the long and short time behaviour of the gel, showing that it is governed by the balance between cell and osmotic forces. The gel equilibrates when the cell and solvent potentials inside the gel are in balance with the external free energy; similarly, the early time behaviour and stability of equilibria is determined by the gradients of these functions inside the gel. Through deriving an analytical solution for the system’s short time behaviour, we investigated how steady states respond to spatial perturbations and determined whether these perturbations will grow or decay.

This analysis also allowed us to evaluate how steady state values and stability change with variations in parameter values. As discussed in Section 2.3, due to the boundary conditions we use, the standard free energy constants 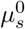 and 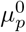 do not appear in the final system of equations. As a result of this, we are not able to reproduce the examples of bistable equilibria presented in Keener *et al*. (2011a) where 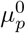 is varied against *θ_p_*. However, as mentioned in Section 2.3, the parameters 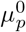 and 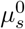 are typically not present in studies using the Flory-Huggins free energy.

In a laboratory setting, this analysis of parameter values and steady state outcomes can be used to predict experimental outcomes given specific gel configurations, *e.g*. suggesting whether a gel will equilibrate or dissolve, or if a different configuration is needed to reach a desired experimental result. This analysis may also allow for physical parameter values to be determined given comparison with experimental results. For example, if we are given experimental data for an initial gel configuration and its subsequent equilibrium state, we may be able to determine that particular parameters must lie within certain ranges through comparison with such steady state diagrams as presented in Fig. 2.

In Section 5, we presented novel results relating to the gel’s evolution, these being the presence of nonuniform equilibria and the case where the polymer fraction and cell density oscillate between increases and decreases. Spatially non-uniform steady states were found to eventuate in the cell-gel system from non-uniform initial conditions in the polymer fraction or cell density in the absence of diffusion. With small diffusion, quasi-steady states were found where the gel evolved to a state with spatial variations present in the variables, but which then gradually smoothed over time. In the oscillating case, due to competition between the free energy and cell traction, we see the fraction of polymer and the cell density repeatedly fluctuate within a spatial region as the gel swells, before it eventually dissolves. To our knowledge, this behaviour has not been described in the literature before; this could be investigated experimentally.

Recent experimental work has suggested using osmotic pressure as a way to impose a desired mechanical compression on cells cultured *in vitro* (Monnier *et al*., 2016; Dolega *et al*., 2017). Our model provides a framework to quantify and evaluate such methods. Although to our knowledge, no one has yet used osmotic pressure to impose dynamic cycles of compression or tension on cells within a gel, our results suggest that this might be possible by, for example, changing the composition of the solution in the solvent bath surrounding the gel as a function of time. Again, this model could be used to predict the ensuing cycles of gel expansion and contraction, as well as to match the frequency and amplitude of these cycles to those seen *in vivo*. This might be beneficial in culturing cartilage cells for example, where oscillating stresses can lead to better mechanical and cell properties in the cells and tissues grown *in vitro* (Salinas *et al*., 2018). Oscillating fluid flow has also been seen to be an important mechanism in areas like proteoglycan production (Eifler *et al*., 2006) and regulating calcium concentrations (Edlich *et al*., 2001).

Our analysis has focussed on the qualitative behaviours predicted by our model in a simple 1D setting. To facilitate greater comparison with experiments, for example that presented in Moon and Tranquillo (1993), a couple of different steps could be taken. An extension to our work here, if a consistent set of data for relevant experiments was available, would be to fit such experimental data for our model parameters and initial conditions, allowing for a more direct comparison between these experiments and simulations like those presented in Section 5. Transforming the model to spherically symmetric coordinates would also help in comparing our results with models looking at spheres of gel like those in Moon and Tranquillo (1993) and Green *et al*. (2013), although we note that, given that such a model would be one-dimensional in the gel’s radius *r*, we would not expect significantly different qualitative outcomes to those seen with our model here, which is one-dimensional in the gel’s length.

In Figs. 8a and 8b, we saw the contraction of a gel with a small initial polymer fraction. In this instance, the polymer fraction and cell density increased gradually at the beginning and end of the gel’s evolution, bookending a period of rapid contraction where *θ_p_* and *n* increased significantly. This behaviour is comparable to examples of mechanically-driven gel contraction presented in Moon and Tranquillo (1993) and Green *et al*. (2013). There was an initial lag in the evolution of the gel’s radius seen in Green’s model which replicates experimental observations from Moon and Tranquillo (1993). This initial lag was not present in Moon and Tranquillo’s mathematical model, and similarly, we did not see an initial lag in changes to the gel’s length in our simulations. Moon and Tranquillo posit that the initial delay seen experimentally is a consequence of the cells spreading after being seeded; it may not be present in our simulations as a consequence of the cells being smoothly distributed at initial time, therefore not requiring a lead time to redistribute themselves through the gel as may happen *in vitro*.

We also saw in Section 5.4 that, with all else held constant, the gel reached a smaller equilibrium length with a smaller initial polymer fraction. Similarly, with cells present, a smaller initial polymer fraction resulted in a larger value for the polymer fraction at equilibrium. Our model therefore captures the negative correlation between the initial polymer concentration and final concentration highlighted in the experimental study presented by Stevenson *et al*. (2010), who also reference this behaviour occurring in experiments such as Zhu *et al*. (2001) and Evans and Barocas (2009).

In the absence of cells, in Fig. 8c and associated simulations, we saw that decreasing the initial polymer fraction *θ_i_* similarly led to gels with a smaller equilibrium length, *i.e*. gels that have contracted further. Without cells, the equilibrium value of the polymer fraction *θ*\* is determined by the parameters in the free energy function; therefore, unlike what was seen for cell-gel systems, *θ*\* remained the same with increases in *θ_i_*. We can therefore see that cell forces play an important role in the negative correlation between initial and final polymer concentrations seen experimentally.

Studies such as Barocas *et al*. (1995) that estimate the value of the cell traction parameter *τ*_0_ typically do so using models that focus on cell-gel mechanics, *i.e*. they do not include the presence of chemical potentials. We have demonstrated herein that chemical potentials can counteract cell traction and affect the degree of gel contraction witnessed. This therefore indicates that without considering the free energy in the gel-solvent system, current models may in fact underestimate the magnitude of cell traction stresses, since the degree of compaction in the experiment will also depend on the mixing energy of the polymer and solvent. It also highlights that the measure of the traction parameter may be quite experiment-specific, depending on the particular configuration of the gel and surrounding fluid. This is supported by Fig. 2d, where we see the balance between cell traction *τ*_0_ and mixing energy *χ* that maintains the same equilibrium value of polymer; increasing *χ* indicates that the gel can equilibrate with a smaller value of *τ*_0_, and vice versa.

We remark that while we have chosen a particular form for the cell force function *G* here, other modelling choices have been used. Green *et al*. (2013) for example numerically investigate numerous different cell force functions: the Murray force function similar to that described in Section 2.2; a ‘preferred ECM density’ function; and functions incorporating chemical concentration. Green *et al*. also consider whether contact inhibition should be incorporated in the form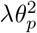 as opposed to the form *λn*^2^, *i.e*. acting on the polymer network instead of the cell density. An extension to this work would be to consider different forms of the cell force function, and again, given consistent experimental data to fit the model, to determine if better agreement is found with particular choices.

In the Moon and Tranquillo (1993) study, the gel is constructed as a microsphere; however, in cell-gel experiments, the gels are often thin layers, where the height of the gel is small relative to its length or radius. Therefore, another extension to this model is to study the gel as a two-dimensional thin film, exploiting the ratio of the film’s vertical and horizontal length scales as a small parameter to rescale the system and derive a reduced system of equations. In this way, gel behaviour in different experimental settings might be compared and analysed. We aim to present such a model in a future publication.

## Acknowledgments

This paper is based on material from JRR’s PhD thesis (Reoch, 2020). JRR acknowledges funding from a University of Adelaide divisional scholarship, and a Westpac STEM PhD scholarship. He is also grateful to the School of Mathematics and Statistics, University of Sydney for providing him with office facilities during a period as a visiting student. JEFG acknowledges funding from an ARC DECRA (DE130100031). All three authors thank Professor Mary Myerscough for helpful discussions.

